# AI aided design of epitope-based vaccine for the induction of cellular immune responses against SARS-CoV-2

**DOI:** 10.1101/2020.08.26.267997

**Authors:** G. Mazzocco, I. Niemiec, A. Myronov, P. Skoczylas, J. Kaczmarczyk, A. Sanecka-Duin, K. Gruba, P. Król, M. Drwal, M. Szczepanik, K. Pyrc, P. Stępniak

## Abstract

The heavy burden imposed by the COVID-19 pandemic on our society triggered the race towards the development of therapies or preventive strategies. Among these, antibodies and vaccines are particularly attractive because of their high specificity, low probability of drug-drug interaction, and potentially long-standing protective effects. While the threat at hand justifies the pace of research, the implementation of therapeutic strategies cannot be exempted from safety considerations. There are several potential adverse events reported after the vaccination or antibody therapy, but two are of utmost importance: antibody-dependent enhancement (ADE) and cytokine storm syndrome (CSS). On the other hand, the depletion or exhaustion of T-cells has been reported to be associated with worse prognosis in COVID-19 patients. This observation suggests a potential role of vaccines eliciting cellular immunity, which might simultaneously limit the risk of ADE and CSS. Such risk was proposed to be associated with FcR-induced activation of proinflammatory macrophages (M1) by Fu et al. 2020 and Iwasaki et al. 2020. All aspects of the newly developed vaccine (including the route of administration, delivery system, and adjuvant selection) may affect its effectiveness and safety. In this work we use a novel in silico approach (based on AI and bioinformatics methods) developed to support the design of epitope-based vaccines. We evaluated the capabilities of our method for predicting the immunogenicity of epitopes. Next, the results of our approach were compared with other vaccine-design strategies reported in the literature. The risk of immuno-toxicity was also assessed. The analysis of epitope conservation among other *Coronaviridae* was carried out in order to facilitate the selection of peptides shared across different SARS-CoV-2 strains and which might be conserved in emerging zootic coronavirus strains. Finally, the potential applicability of the selected epitopes for the development of a vaccine eliciting cellular immunity for COVID-19 was discussed, highlighting the benefits and challenges of such an approach.

## 1 Introduction

As of August 6, 2020, more than 19 million cases of COVID-19 were reported worldwide, leading to more than 700 thousands deaths (https://coronavirus.jhu.edu/map.html). The disease was first recorded on December 26, 2019, when a 41-year-old patient with no history of hepatitis, tuberculosis, or diabetes was hospitalized at the Central Hospital of Wuhan due to respiratory problems (F. Wu et al. 2020). The metagenomic RNA sequencing of bronchoalveolar lavage (BAL) fluid sample obtained from that patient led to the identification of the seventh coronavirus (CoV) strain known to infect humans.

Coronaviruses are well known human respiratory pathogens associated with the common cold. Until the 21st century they were neglected by the medical world, but the emergence and subsequent spread of the SARS-CoV in the 2002/2003 season raised interest in this virus family and increased awareness of the potential threat. At present, there are four seasonal coronaviruses infecting humans and they cluster within alphacoronaviruses (HCoV-NL63, HCoV-229E) and betacoronaviruses (HCoV-OC43, HCoV-HKU1) genera. Further, three zoonotic strains were reported - severe acute respiratory syndrome coronavirus (SARS-CoV; 2002-2003), the Middle East respiratory syndrome coronavirus (MERS-CoV; 2012-), and SARS-CoV-2 (2019-), all of which belong to the betacoronavirus genus (A. Wu et al. 2020). The highly pathogenic species cluster in two subgenera – sarbecoviruses (SARS-CoVs) and merbecoviruses (MERS-CoVs) (Hu et al. 2018; F. Wu et al. 2020; Zhou et al. 2020).

While generally, viruses infect one host, some have broader specificity or can cross the interspecies borders, causing outbreaks, epidemics, and pandemics. In this context, it is worth mentioning viruses like the Ebola virus, dengue fever virus, Nipah virus, rabies virus, or Hendra virus. However, these are well known and long studied animal viruses that only sometimes enter the human population. Coronaviruses are slightly different, as among the myriads of viral species and subspecies found in animals, it is unlikely to predict the place, the time, and the genotype of the coronavirus that will emerge. The classic transmission route of these viruses encompasses the spillover of the bat species to wild or domesticated animals, rapid evolution in this intermediate host, and subsequent transmission to humans. Coronaviruses emerge at different sites of the globe where the interaction between humans and animals is broad, such as the Asian wet markets and the dromedary camel farms in the Arabian peninsula. While these high-risk regions were identified, the next spillover may occur in Europe or the Americas, as sarbecoviruses are prevalent around the globe (Andersen et al. 2020).

The coronaviral genome is a single-stranded RNA of positive polarity, which ranges in size from 26,000 up to 32,000 bases. Two-thirds of the genome on the 5’ end are occupied by two large open reading frames (ORFs) that may be read along due to the ribosomal slippery site. The resulting polyprotein undergoes subsequent autoproteolysis, and the matured proteins form the complete replicatory machinery and re-shape the microenvironment of the infection. Downstream of the 1ab ORFs, a number of ORFs are found that encode structural and accessory proteins (Cui et al. 2019; Song et al. 2019). Four major structural proteins are: spike surface glycoprotein (S), envelope protein (E), membrane glycoprotein (M), and nucleocapsid phosphoprotein (N). Of them the S protein is the primary determinant of the species and cell tropism, interacting with the receptors and co-receptors on the host cells (Li 2016; Zhu et al. 2020).

Evolutionary studies indicate that CoV genomes display high plasticity in terms of gene content and recombination (Forni et al. 2016). The long CoV genome expands the sequence space available for adaptive mutations, and the spike glycoprotein used by the virus to engage target cells can adapt with relative ease to exploit homologs of cellular receptors in different species. While coronaviruses are rapidly evolving, their mutation rate is lower than expected for an RNA virus. The large genomes require proofreading machinery to maintain their functions, and proteins required for such activity are among the 1a/1ab proteins.

While sarbecoviruses and merbecoviruses are associated with severe, potentially lethal diseases and are known for their epidemic potential in humans and animals, several years of research did not allow for the development of effective and safe vaccines. In addition to the high variability and ability to elude immune recognition, there are several aspects to be considered. First, the antibody-dependent enhancement (ADE) of the infection was previously reported for some coronaviruses, including sarbecoviruses. ADE is based on the fact that the virus exploits non-neutralizing antibodies to enter the host’s cells utilizing the Fc receptor (FcR). The ADE phenomenon was originally observed for antibodies specific to certain dengue virus serotypes developed after a primary infection. During subsequent dengue infections, caused by a different virus serotype, these antibodies were able to recognize the virus but were not capable of neutralizing it. Instead, antibodies bridged the dengue virus and the Fc receptors of the immune cells, such as macrophages and B-cells, mediating viral entry into these cells and transforming the disease from a relatively mild illness to a life-threatening infection. A similar mechanism was later observed for HIV and Ebola infections (Beck et al. 2008; Dejnirattisai et al. 2010; Guzman et al. 2007; Katzelnick et al. 2017; Takada et al. 2003; 2001; Whitehead et al. 2007; Willey et al. 2011). Importantly, ADE has also been reported for some coronaviruses. The best-documented ADE cases are associated with feline infectious peritonitis virus. It was shown that immunization of cats with feline coronavirus spike protein leads to increased severity during future infections due to the induction of infection-enhancing antibodies (Corapi et al. 1992; Hohdatsu et al. 1998). Further, some studies show that antibodies induced by the SARS-CoV spike protein enhance viral entry into FcR-expressing cells (Jaume et al. 2011; Kam et al. 2007; S.-F. Wang et al. 2004). It was confirmed that this Abs-dependent SARS-CoV entry was independent of the classical ACE2 receptor-mediated entry (Jaume et al. 2011). A recent study investigated the molecular mechanism behind antibody-dependent and receptor-dependent viral entry of MARS-CoV and SARS-CoV pseudoviruses in vitro (Y. Wan et al. 2019). The authors demonstrated that MERS-CoV and SARS-CoV neutralizing monoclonal antibodies (mAbs) binding to the receptor-binding domain region of the respective spike protein were capable of mediating viral entry into FcR-expressing human cells, confirming the possibility of coronavirus-mediated ADE. Given the critical role of antibodies in host immunity, ADE causes serious concerns in epidemiology, vaccine design, and antibody-based drug therapy.

The consequences of ADE may be dramatic, as it may cause lymphopenia and induce or increase the frequency of the cytokine storm syndrome (CSS). This may result directly from the active infection of immune cells, which in response produce large amounts of the inflammatory markers or indirectly, when virus-antibody complex binds to FcR and activates pro-inflammatory signaling, skewing macrophages responses to the accumulation of pro-inflammatory M1 macrophages in lungs. The macrophages secrete inflammatory cytokines, such as MCP-1 and IL-8, which lead to worsened lung injury (Fu et al. 2020). In both animal models and patients who eventually died from SARS, extensive lung damage was associated with high initial viral loads, increased accumulation of inflammatory monocytes/macrophages in the lungs, and elevated levels of serum pro-inflammatory cytokines and chemokines (IL-1, IL-6, IL-8, CXCL-10, and MCP1) (Channappanavar et al. 2016). Moreover, during the SARS-CoV outbreak in Hong Kong (2003-2004), 80% of the patients developed acute respiratory distress syndrome after 12 days from the diagnosis, coinciding with IgG seroconversion (Peiris et al. 2003). Another study by Huang et al. 2020 highlighted an increased release of IL-1β, IL-4, IL-10, IFNγ, MCP-1, and IP-10 in COVID-19 patients. Interestingly, compared with non-severe cases, severe patients in the intensive care unit showed higher plasma concentrations of TNFα, IL-2, IL-7, IL-10, MIP-1A, MCP-1, and G-CSF, supporting the hypothesis of a possible correlation between CSS and severity of the disease. An extensive study done by Liu et al. 2019 demonstrated that anti-spike IgGs enhanced the induction of pro-inflammatory cytokines (i.e., IL-6, IL-8, and MPC-1) in Chinese rhesus monkeys through the stimulation of alternatively activated monocyte-derived macrophages (MDM) upon SARS-CoV rechallenge. The presence of high MDM infiltrations was shown by histochemical staining of the lung tissue from 3 deceased SARS patients. The blockade of Fc-receptors for IgG (FcγRs) reduced proinflammatory cytokine production, suggesting a potential role of FcγRs for the reprogramming of alternatively activated macrophages. Putting these results in the context of other works in literature (Pahl et al. 2014), one has to consider that anti-S IgG may promote pro-inflammatory cytokine production and, consequently, CSS development.

Taking into account the risk associated with the improper humoral response and high variability of sites targeted by the neutralizing antibodies, together with the low effectiveness of IgG-mediated immunity during mucosal infection, it is of importance to consider the anticoronaviral vaccine in a broader perspective. This may include alternative delivery systems/routes based on, e.g., virus-like particles and intranasal delivery for the IgA mediated response, but it is also important to consider combining the humoral response with the cell-mediated response. Ideally, such an approach might allow for the design of a vaccine carrying carefully selected epitopes to induce only the neutralizing antibodies and epitopes targeted for induction of the cellular response. While neutralizing antibodies impair the virus entry, activated CD8+ cytotoxic T-cells can identify and eliminate infected cells. Moreover, CD4+ helper T-cells are required to stimulate the production of antibodies. Antibody response was found to be short-lived in convalescent SARS-CoV patients (Tang et al. 2011) in contrast to T-cell responses, which have been shown to provide long-term protection (Fan et al. 2009; Peng et al. 2006; Tang et al. 2011), up to 11 years post-infection (O.-W. Ng et al. 2016). The activation of CD8+ cytotoxic T-cells capable of recognizing and destroying infected cells represents a crucial second line of defense against the virus that should be considered. The importance of both CD8+ and CD4+ T-cell activation has been reported in several SARS-CoV studies for both animal models and humans (Channappanavar et al. 2014). Moreover, several recent studies indicate a strong correlation between the reduction of lymphocyte counts (CD4+ and CD8+) and the severity of COVID-19 cases (N. Chen et al. 2020; Liao et al. 2020; S. Wan et al. 2020).

The selection of epitopes capable of eliciting either B-cell or T-cell responses is a critical step for the development of subunit vaccines. Most of the efforts in this area are directed towards the stimulation of neutralizing antibodies, whereas the cellular immune response is less explored. Considering the importance of T-cell activation for vaccine efficacy, the focus of the work here presented is on the latter. Despite the apparent similarity between SARS-CoV and SARS-CoV-2, there is still a considerable genetic variation between these two. Thus, it is not trivial to assess if epitopes eliciting an immune response against previous coronaviruses are likely to be effective against SARS-CoV-2, with the exception of identical peptides shared among subgenera. A restricted list of SARS-CoV epitopes identical to those present in SARS-CoV-2 and resulting positive in immunoassays, has been recently reported (Ahmed et al. 2020). Nonetheless, the 29 T-cell epitopes described therein are mostly limited to S, N, and M antigens and encompass an exiguous number of Class I Human Leukocyte Antigen (HLA) alleles. In order to extend the search area to other epitopes, computational predictive models might be applied. Methods for the selection of vaccine peptides are typically based on the predicted binding affinity (or probability of presentation on the cell surface) of peptide-HLA (pHLA) complexes or defined by the physicochemical properties of the peptides (Baruah and Bose 2020; Grifoni et al. 2020; Lee and Koohy 2020). These methods take into account only restricted parts of processes contributing to the final immunogenicity of an epitope, and thus their prediction capabilities are limited. In addition to pHLA binding, proteasome cleavage, pHLA loading, and presentation, as well as direct activation of CD8+ T-cell to the pHLA complex should be taken into account.

Here, we use a machine learning model for the prediction of epitope immunogenicity. The model is trained on data including the experimental T-cell immunogenicity data of viral epitopes. We validate our model on publicly available immunogenicity data of epitopes from the *Coronaviridae* virus family (held out from training). Assessment of the risk of immuno-toxicity and the analysis of epitope conservation among different strains are also performed.

## 2 Materials and Methods

### 2.1 Presentation data

A curated dataset containing peptides presented by class I HLAs on the surface of host cells was extracted from publicly available databases (Abelin et al. 2017; Di Marco et al. 2017; Sarkizova et al. 2020). The presentation of each peptide within the dataset was experimentally confirmed by mass-spectroscopy experiments. All peptides were of human origin and were presented on the surfaces of monoallelic human cell lines (see Figure 1 and Table 1). Synthetic negative data (non-presented peptides) were also prepared based on human proteome (GRChg38, release 98).

**Table 1.** The total number of pHLAs included in our model from each dataset.

| Source publication | No. pHLAs |
| --- | --- |
| Abelin et al. [45] | 22999 |
| Di Marco et al. [46] | 22889 |
| Sarkizova et al. [47] | 146739 |

**Figure 1.**
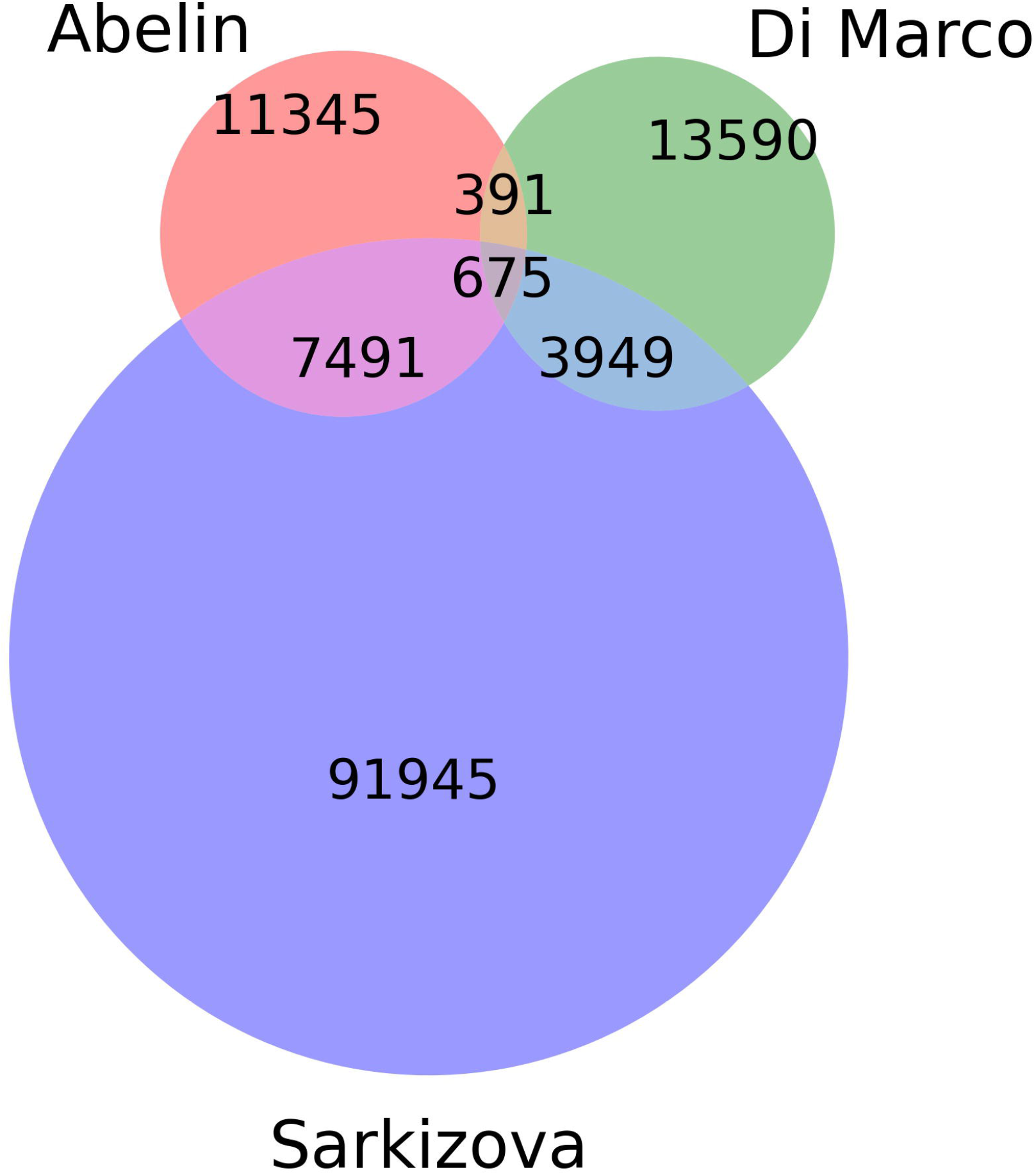
Venn diagram showing the number of unique and common peptides among datasets.

### 2.2 Immunogenicity data

All peptides collected from the IEDB database (Vita et al. 2019) were of viral origin and were confirmed in experimental immunoassays. Similar data were extracted from selected publications (Chen et al. 2005; Liu et al. 2010; Ogishi and Yotsuyanagi 2019; Tsao et al. 2006; Y.-D. Wang et al. 2004; Zhang 2013). The number of pHLAs (per immunoassay category) used for training is given in Table 2. Most of the peptides were obtained from human hosts, with a minority obtained from transgenic mice. Only peptides containing 8-11 amino acids were included in the analysis. In some cases, multiple experimental settings and protocols were used to validate immunogenicity for a given pHLA, occasionally leading to non-consensual results. Each pHLA was considered immunogenic if at least one experiment conducted on human cells positively confirmed that immunological event. If no experiments conducted on human cells were available, the pHLA was considered immunogenic, if at least one such confirming experiment was conducted in transgenic mice. The remaining pHLAs were used as negative examples. From this dataset we held out the *Coronaviridae* family as a separate test set.

**Table 2.** The number of pHLA complexes used for training per immunogenic assay group.

| Source publication | Negative | Positive |
| --- | --- | --- |
| IFNγ | 23249 | 2598 |
| cytotoxicity | 218 | 524 |
| proliferation | 7 | 34 |
| cytokines/chemokines | 0 | 13 |
| TNFα | 1 | 8 |

**Table 3.**
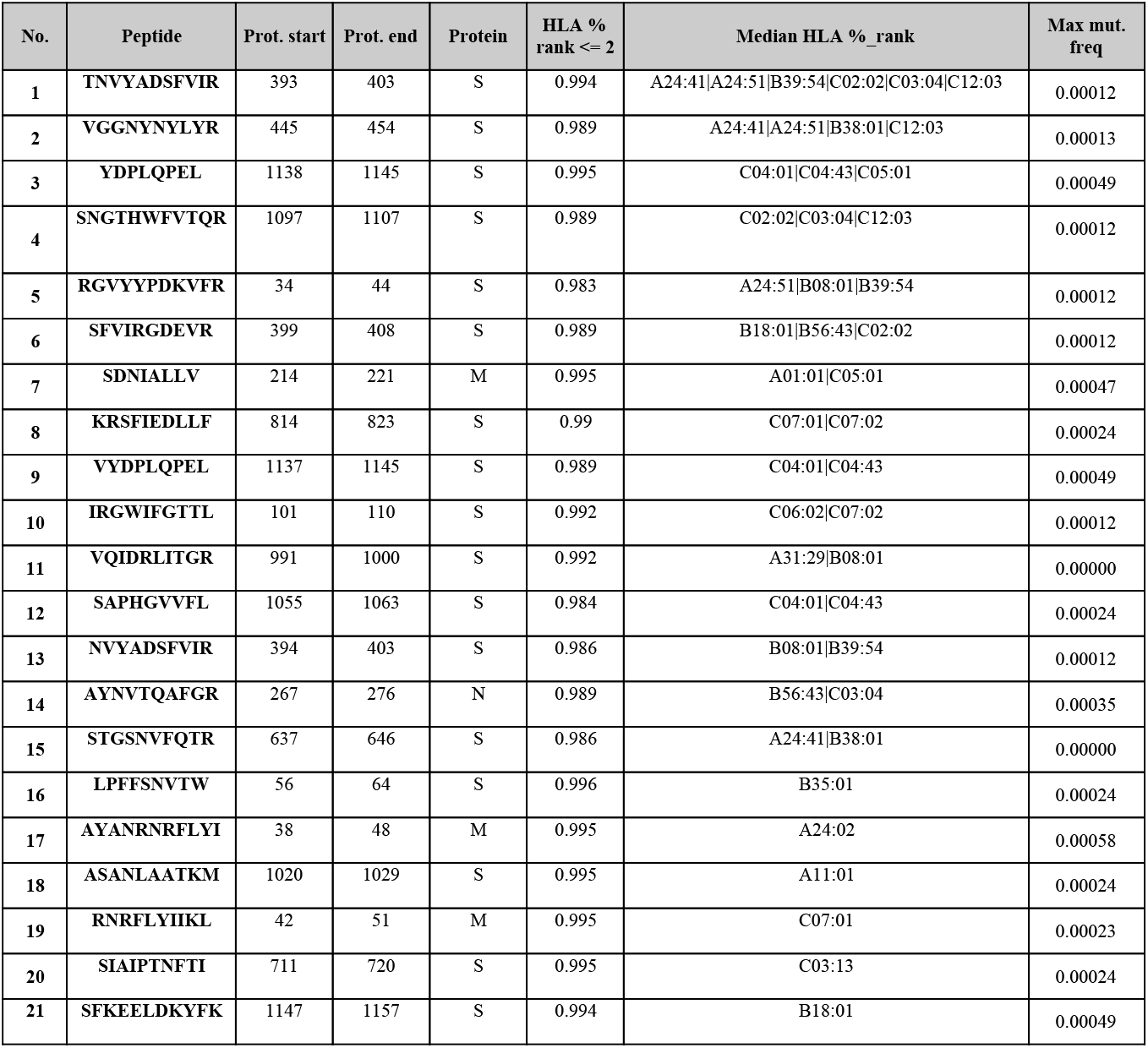
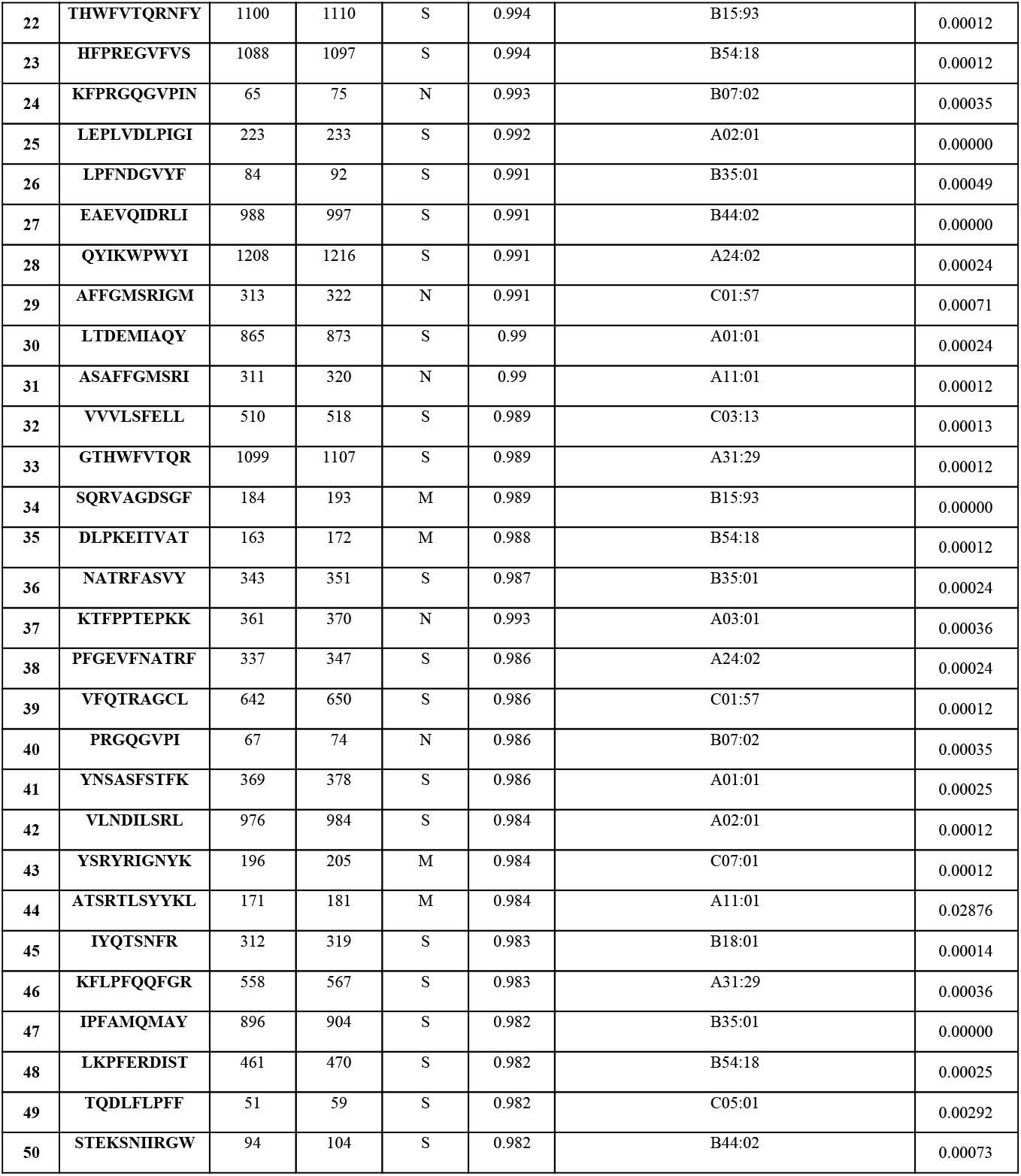
Peptides with ArdImmune Rank percentage rank ≤ 2 obtained from SARS-CoV-2 structural proteins, sorted by (1) the number of HLA types capable of binding and presenting given peptide and (2) the median rank across different HLA types. Peptides marked in red are considered as Highly Variable (HV) due to maximum mutation frequency score >= 0.05

### 2.3 Predictive model design

Our computational methods are based on machine learning and predict (1) the probability of pHLAs to be presented on the host’s cell surface and (2) the immunogenicity of such complexes. The model for pHLA presentation is based on artificial neural networks and has been trained on a curated collection of peptide presentation data (Abelin et al. 2017; Di Marco et al. 2017; Sarkizova et al. 2020). Both peptide sequence and HLA type were taken into consideration as separate inputs. We use bootstrapping and select 80% of positive examples during training with the remaining ones used for early stopping. We then ensemble the results of a collection of 27 such neural networks. Our model is pan-specific and can be used to generate predictions for any peptide and any canonical class I HLA (i.e., A, B, C). Note, that the accuracy of our method depends on the considered HLA type, as in the case of other machine learning methods for predicting pHLA properties.

The model mentioned above was also used as a starting point for training the immunogenicity model. The latter was fine-tuned using the viral peptide immunogenicity data collected from IEDB (Vita et al. 2019) and Ogishi and Yotsuyanagi 2019. The immunogenicity model was validated using a Leave One Group Out (LOGO) cross-validation scheme with groups defined by viral families. The final model is an ensemble of 11 models - one per each LOGO split. An additional group “others” was defined by aggregating data from viruses that belong to several families, having a small number of observations. Such an approach provides data splits according to the virus families and leads to a better measure of performance on virus families not seen in training (e.g., *Coronaviridae*). Moreover, it reveals the differences in model performance on various virus families. The final predictions of our model (called ArdImmune Rank) are obtained by combining the predictions of both models (i.e. the pHLA presentation and the immunogenicity model).

### 2.4 Validation scheme

In order to validate the ArdImmune Rank model over different virus families not seen during the training procedure, a LOGO strategy was applied (note that in this LOGO validation - in Figures 4 and 5 - we use a single immunogenicity model instead of 11 models, as in Figure 3). The peptides associated with coronaviruses were held out from the dataset and left for testing purposes only. At each LOGO iteration, the dataset was split into training and validation sets, and the model was tested accordingly. Peptides within the training set highly similar to the ones in the validation set were removed from the training set. The similarity of peptides was assessed using a clustering algorithm classifying their sequences into groups of peptides sharing a common root (differing only by short prefixes or suffixes of lengths of at most 3 amino acids). The number of pre-processed peptides in each group is given in Figure 2. Finally, the immunogenicity model (an ensemble of 11 models from the LOGO scheme) was validated on the held-out *Coronaviridae* dataset.

**Figure 2.**
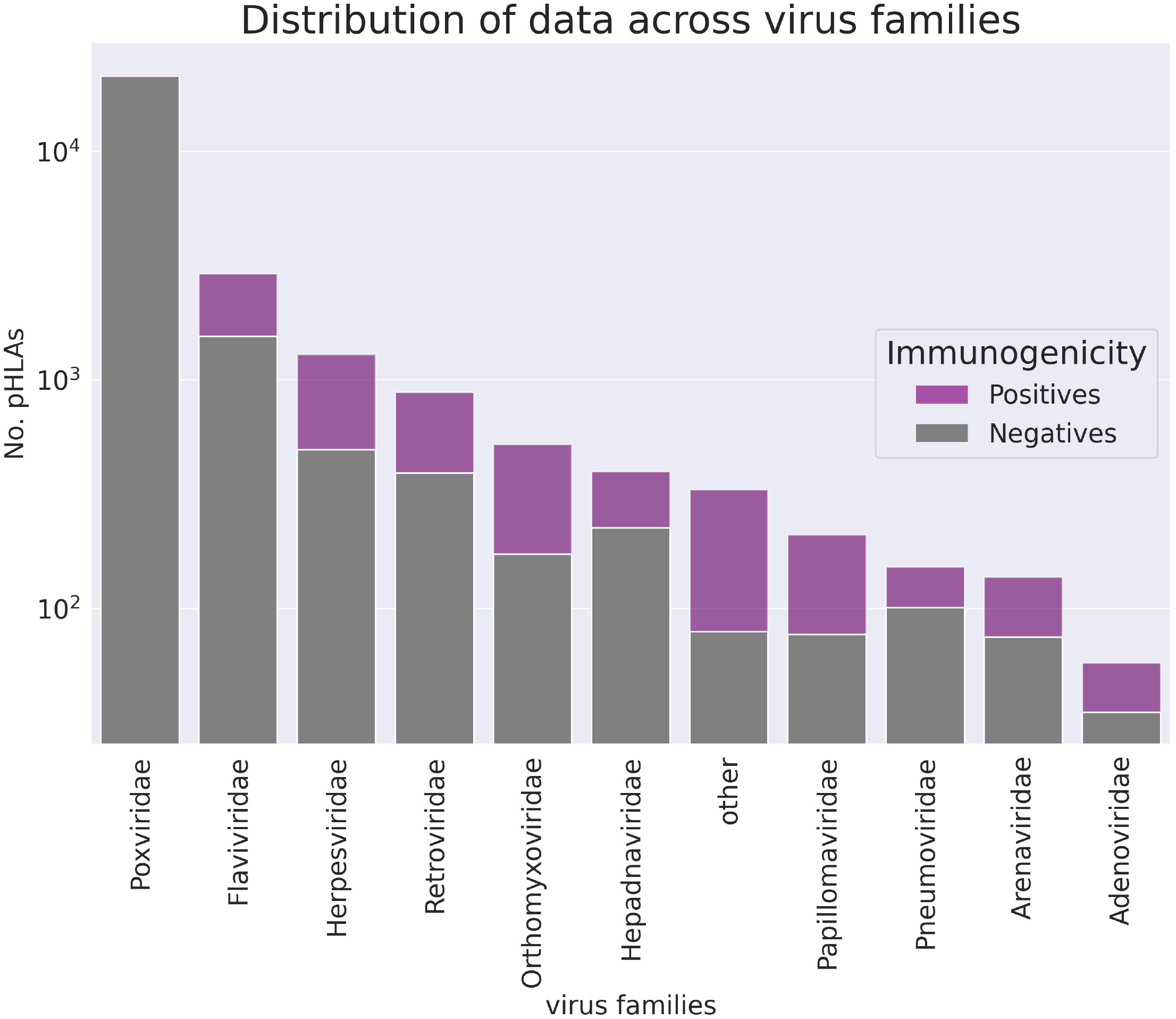
The number of pHLA complexes with confirmed immunogenicity in the curated database per virus family (logarithmic scale). Families counting less than 55 observations are aggregated in the “other” group.

**Figure. 3.**
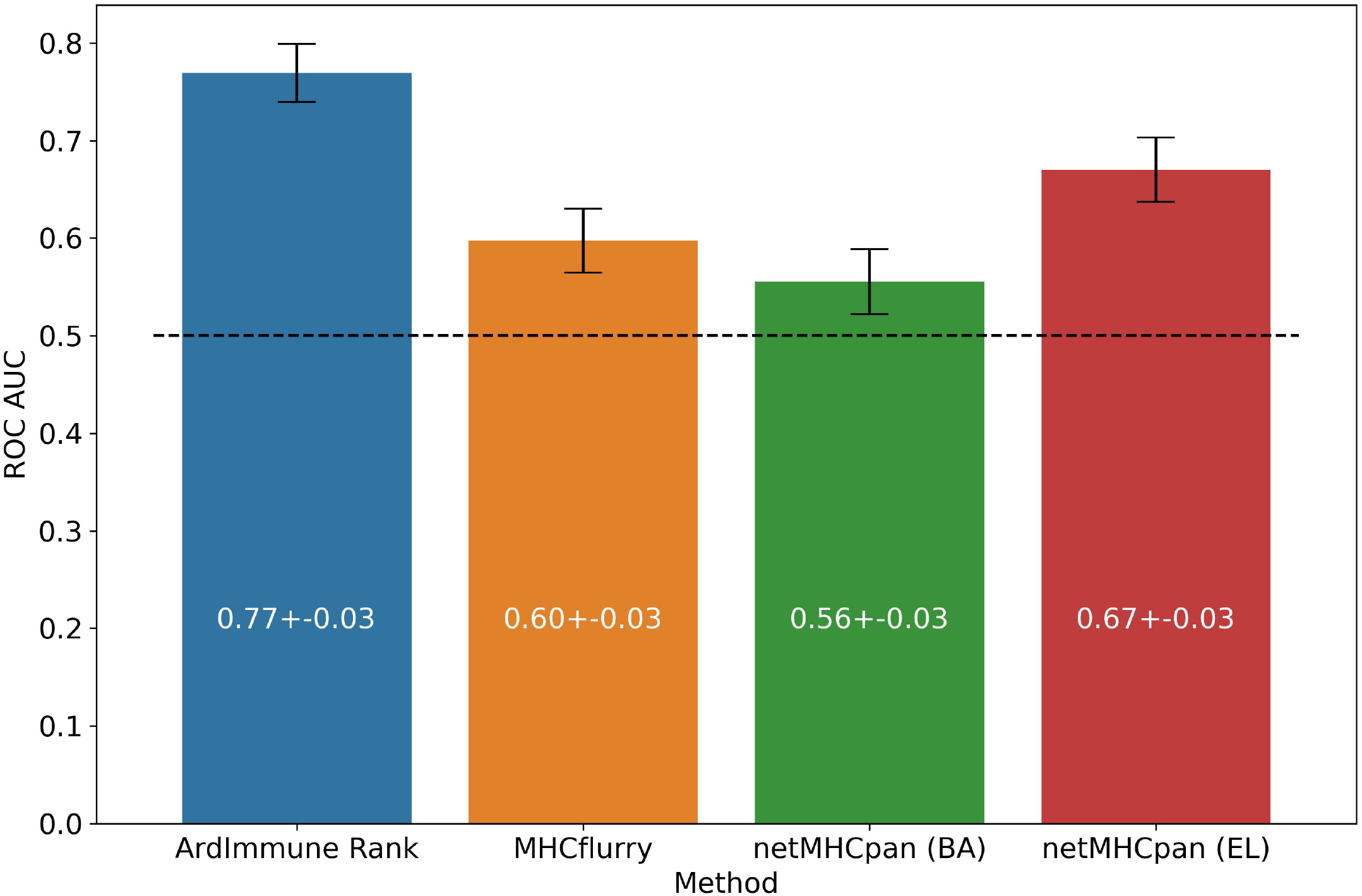
Predictive performance of the selected models on the *Coronaviridae* dataset. ArdImmune Rank: blue bars, MHCflurry: brown bars. netMHCpan: green and red bars for the predicted binding affinity (BA) and ligand likelihood (EL), respectively.

**Figure 4.**
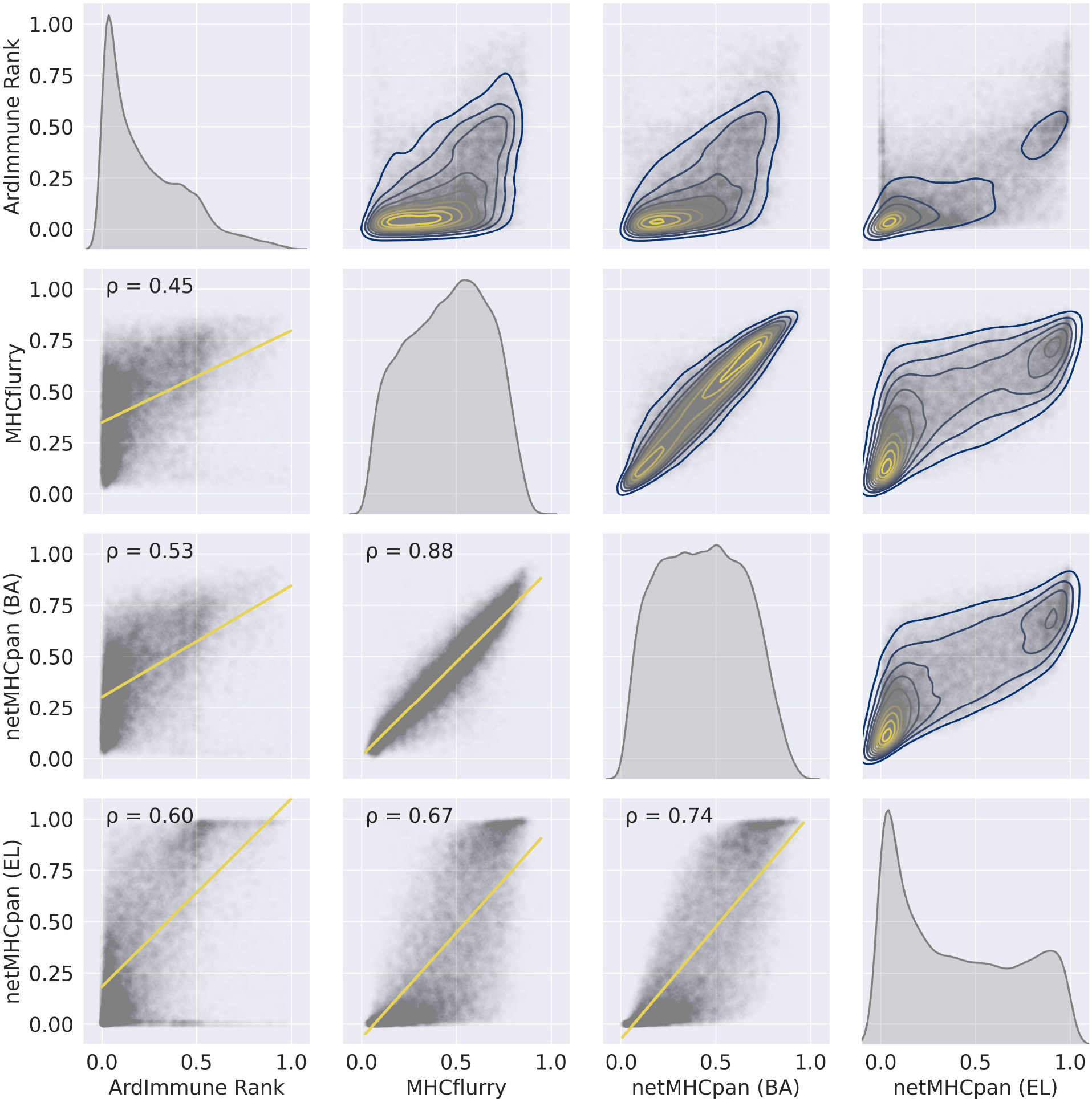
The pairwise relationships between the predictions of the selected models on the training set - (1) ArdImmune Rank, (2) MHCflurry, (3) netMHCpan (BA), and (4) netMHCpan (EL). Lower triangle - scatterplots with linear regression models fitted (yellow lines) and Pearson’s correlation coefficients (PCC) that measure linear correlations between two variables. Diagonal and upper triangle - the prediction distributions obtained by kernel density estimations (1D-KDE and 2D-KDE respectively).

**Figure 5.**
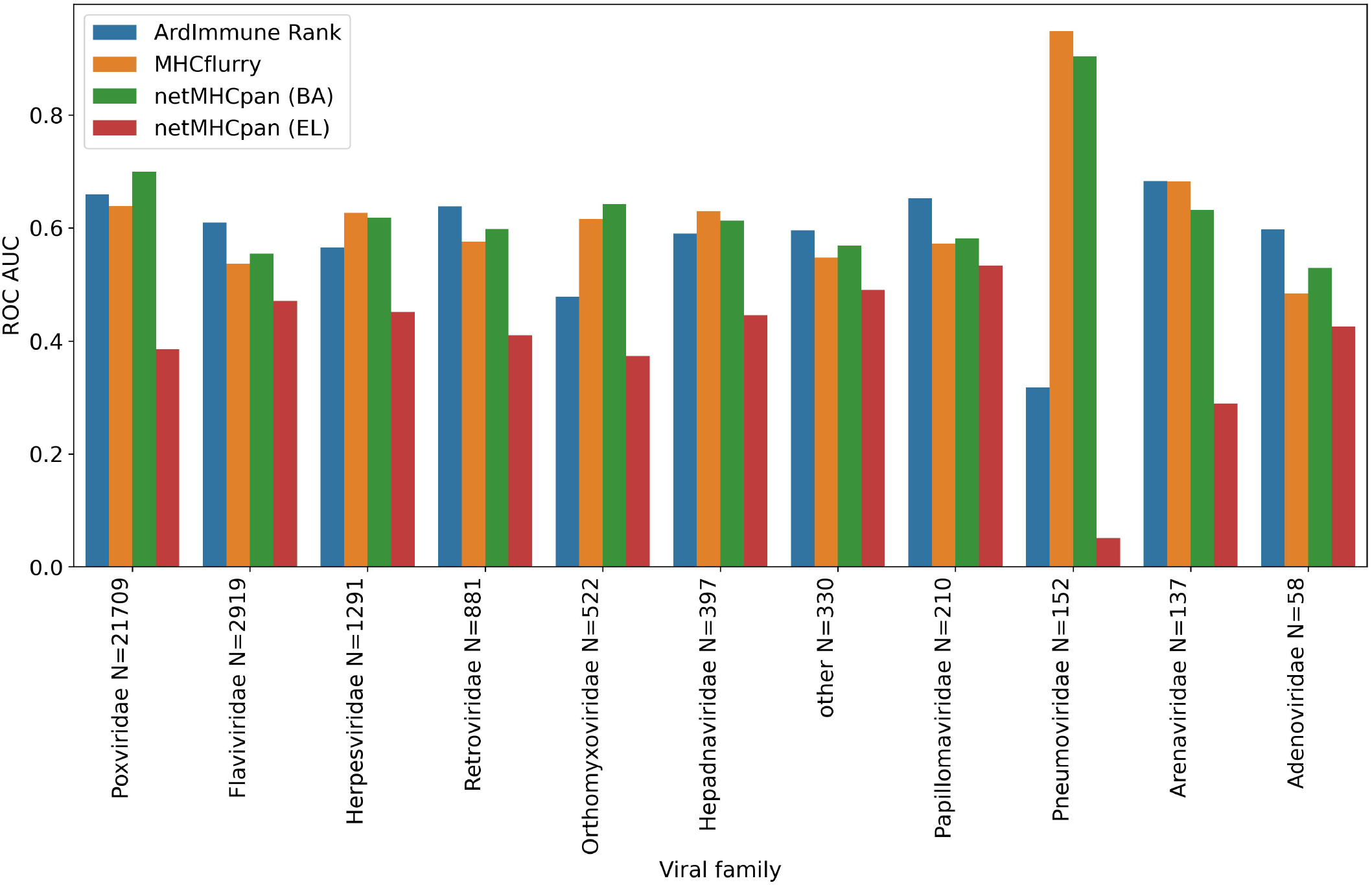
Predictive performance of the selected models obtained in a LOGO cross validation and measured with ROC AUC. ArdImmune Rank: blue bars, MHCFlurry: orange bars, NetMHCpan (BA): green bars, NetMHCpan (EL): red bars.

### 2.5 SARS-CoV-2 data analysis

#### 2.5.1 Selection of HLA alleles

Class I HLA types were chosen based on their frequency of occurrence in the USA and Europe. HLA-allele frequency data were downloaded from http://www.allelefrequencies.net/, accounting for all the populations within the regions of choice and all ethnicities. The overall frequency for each allele was computed as the weighted average with weights corresponding to the size of each population, separately for the USA and Europe, encompassing all ethnic populations. All HLA-alleles with frequency ≥ 0.01 were chosen for the study.

#### 2.5.2 Toxicity/tolerance evaluation

In order to evaluate the risk for a given pHLA to be cross-reactive or tolerogenic with respect to self-epitopes within the human proteome, a procedure for the evaluation of potential toxicity/tolerance was implemented. Initially, each SARS-CoV-2 peptide was queried against the reference human proteome (GRChg38, release 100) using the BLASTp algorithm and a BLOSUM45 substitution matrix. All matches with e-values less than or equal to 4 were included in the analysis. The selected peptides are available in Supplementary Data 1.

#### 2.5.3 Selection of peptides

The dataset consisting of SARS-CoV-2 peptides was generated according to the following procedure: (1) all the reference sequences of the virus proteins were collected from the NCBI database (https://www.ncbi.nlm.nih.gov/search/all/?term=SARS-CoV-2), (2) from each protein, all possible peptides of length 8-11 amino acids were selected. In addition, for proteins encoded by the ORF1a and ORF1ab genes (i.e., pp1a, pp1ab, respectively), the peptides within the cleavage sites were excluded. Finally, all the peptide duplicates were removed from the dataset. A total of 47,612 peptide sequences were collected.

#### 2.5.4 Estimation of SARS-CoV-2 genome diversity

The analysis of conservation of SARS-CoV-2 genomic sequences was performed using 8,639 complete genomic sequences obtained from the GISAID database (https://bigd.big.ac.cn/ncov/release_genome) and GenBank (https://www.ncbi.nlm.nih.gov/genbank/sars-cov-2-seqs/). All sequences were aligned to the SARS-CoV-2 reference genome (NCBI Reference Sequence: NC_045512.2). The R DECIPHER package (Wright 2015) was used for the multiple sequence alignment (MSA). Next, all nucleotides within the coding cDNA sequence (CDS) regions of the reference genome were translated into proteins using the R Biostring package (Pagès et al. 2020). All the fuzzy codons were marked as unknown amino acids. For each protein, all sequences containing indels or being inconveniently aligned were removed. Mutation frequencies were computed for each amino acid in the SARS-CoV-2 proteome. The mutation frequency of each amino acid was defined as the ratio between the number of translated protein sequences containing the mutation and the number of sequences containing a valid nucleotide (sequences containing unknown nucleotides in this position were excluded). The maximum mutation frequency score for each peptide was computed as the maximum value of the mutation frequency scores among all amino acid positions of the peptide. Mutation frequency values for all positions within SARS-CoV-2 proteome are available in Supplementary Data 2.

### 2.6 Datasets for external comparison

In order to highlight similarities and differences of our approach with respect to other methods, we compare the scores of our model with scores relative to the same pHLAs reported in a list of selected studies. A peptide missing from the reference proteome (“QSADAQSFLNR”) was removed. Only peptides between 8 and 11 amino acids were considered. The peptides arising from the cleavage sites of the ORF1a/ab polyprotein were also removed from the datasets. This procedure led to the following datasets:

1. Baruah and Bose 2020: 5 epitopes from the surface glycoprotein of SARS-CoV-2 and their corresponding HLA class I supertype representative were reported by the authors (Table 1 in the reference publication). Bioinformatics protocols, machine learning methods, and structural analysis were applied in the original paper for the selection of these pHLAs.
2. Lee and Koohy 2020: 19 A*02:01 restricted epitopes were selected applying TCR-specific Position Weight Matrices (PWM) previously published by the authors. The geometric mean of the three scores was used as an estimator for immunogenicity (Tables 4 and 5 in the reference publication).
3. Grifoni et al. 2020:
  a. 1st dataset: 386 SARS-CoV-2 CD8+ predicted epitopes were collected (Table S6 in the reference publication) and 41 peptides were excluded as a result of our filtering procedure.
  b. 2nd dataset: 28 SARS-CoV-2 CD8+ epitopes mapped to immunodominant SARS-CoV epitopes were selected (Table 5 in the reference publication). One peptide was excluded as a result of our filtering procedure.
4. Gupta et al. 2020: 10 A*11:01 restricted peptides from the surface glycoprotein of SARS-CoV-2 were selected by the authors (Table 4a in the reference publication). Bioinformatics protocols, machine learning methods, and structural analysis were used for the selection of those pHLAs. A candidate with an optimal docking score is reported.
5. Prahar et al. 2020: 138 peptides with pHLA complex stability measurements performed using Immunotrack’s NeoScreen® assay were made available by the authors. A peptide absent in our dataset was excluded from the comparison.
6. Rammensee et al. 2020: 5 HLA class I peptides were used by the authors for the experimental vaccination of self-experimenting healthy volunteers. IFNγ ELISPOT assays for the measurement of CD8+ activation were negative for all these peptides.
7. Smith et al. 2020: Predictions for ∼615k peptides were extracted from the supplementary table S1 of the reference publication. Approximately 7600 peptides were excluded as a result of our filtering procedure.

The ArdImmune Rank percentile rank for the pHLAs described in the above datasets was computed for groups of peptides according to their HLA allele. Only pHLAs with a binding affinity percentile rank score < 0.02 (predicted using NetMHCpan 4.0) were considered. The predictions were calculated separately for peptides of structural and non-structural origin.

**Table 4.**
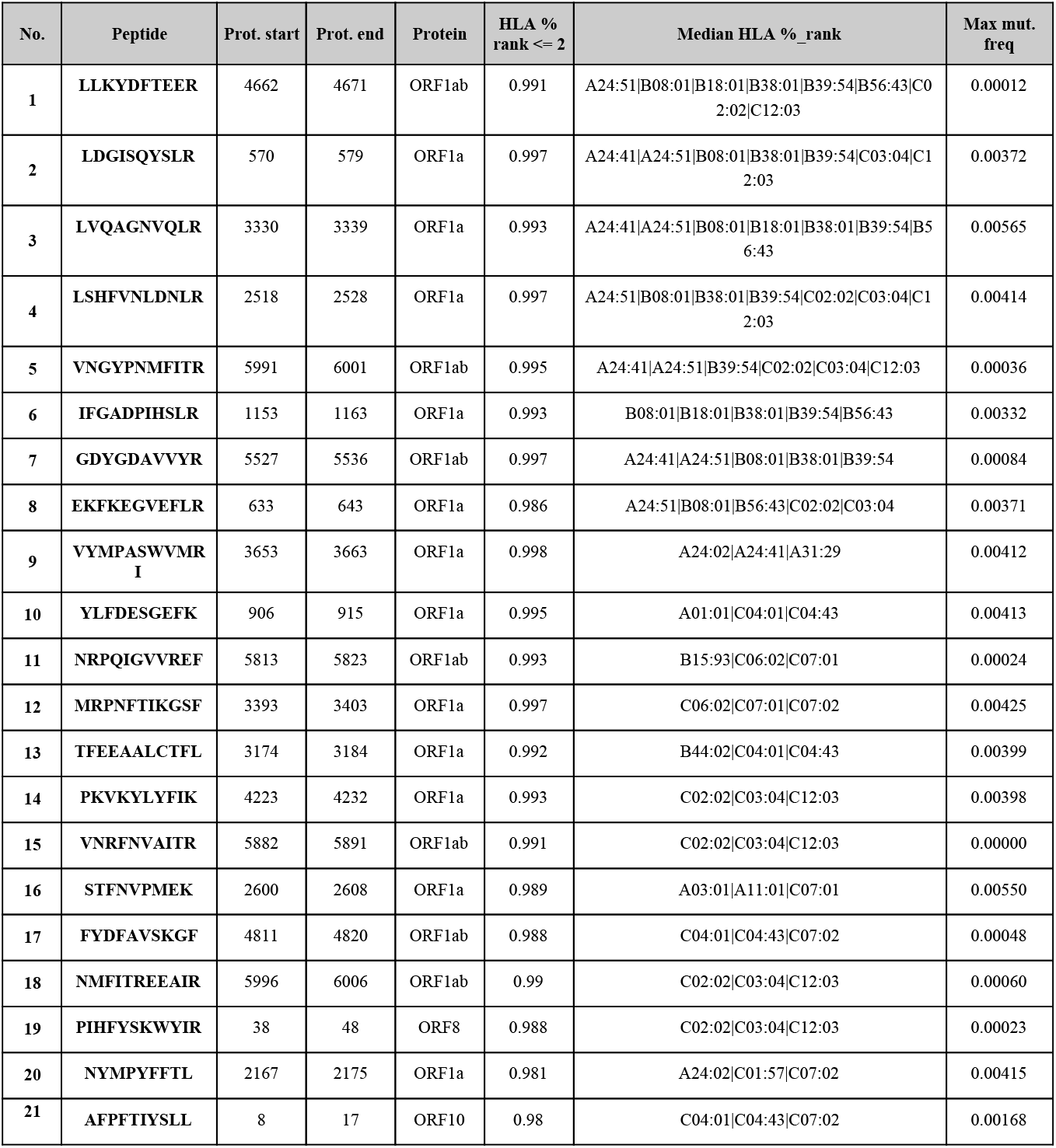
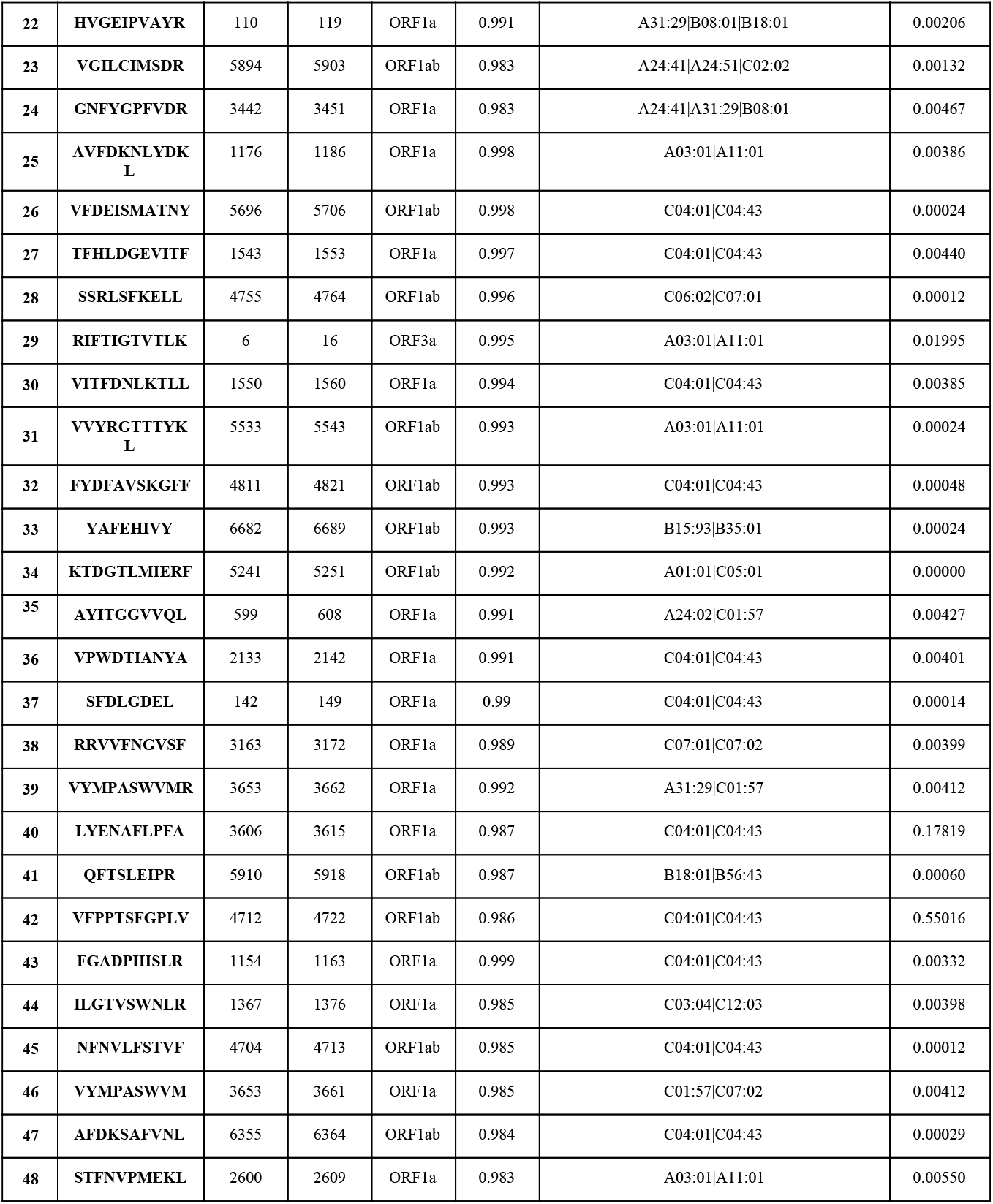
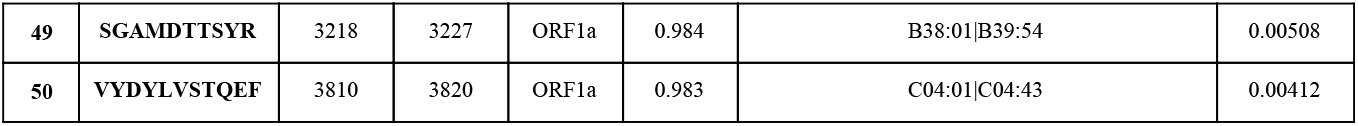
Peptides with model percentage rank ≤ 2 obtained from SARS-CoV-2 non-structural proteins, sorted by (1) the number of HLA types capable of binding and presenting given peptide and (2) the median rank across different HLA types. Peptides marked in red are considered as Highly Variable (HV) due to maximum mutation frequency score >= 0.05

**Table 5.** The most frequently mutated positions within the SARS-CoV-2 proteome.

| No. | Protein | Protein Position | Mutation Frequency |
| --- | --- | --- | --- |
| 1 | ORF1ab | 4715 | 0.5502 |
| 2 | S | 614 | 0.5478 |
| 3 | ORF3a | 57 | 0.1789 |
| 4 | ORF1a | 3606 | 0.1781 |
| 5 | N | 203 | 0.1770 |
| 6 | N | 204 | 0.1765 |
| 7 | ORF1a | 265 | 0.1646 |
| 8 | ORF3a | 251 | 0.1439 |
| 9 | ORF8 | 84 | 0.1384 |
| 10 | ORF1ab | 5865 | 0.0926 |
| 11 | ORF1ab | 5828 | 0.0924 |
| 12 | ORF1a | 765 | 0.0668 |
| 13 | ORF1a | 739 | 0.0590 |

## 3 Results

### 3.1 Model performance

The performance of our method on the test set encompassing *Coronaviridae* epitopes (excl. SARS-CoV-2) is shown in Figure 3. In addition, the results of our approach are compared to those obtained by other commonly used pHLA binding affinity and pHLA presentation probability predictors, namely netMHCpan 4.0 (Jurtz et al. 2017) and MHCflurry (O’Donnell et al. 2018). For both tools MHCflurry and netMHCpan (BA), the binding affinity predictions in nanomoles (nM) are converted into [0, 1] range with a widely used logarithmic transformation (i.e. first the predictions are bounded from above by 50,000 nM and from below by 1 nM and then transformed with 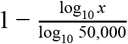). The difference in the predictive performance (measured with ROC AUC) of our model with respect to the other methods is statistically significant (and ranges from 0.10 to 0.21). Moreover, the high Pearson correlation between the results produced by the binding predictors (corr. coeff. ρ=0.88) and the low correlation of such results with the predictions of our model (ρ=0.45 and ρ=0.53) demarcate substantial differences between our approach and the approaches based on those methods for predicting immunogenic epitopes (see Figure 4).

We apply the LOGO cross-validation scheme according to the procedure described in the Materials and methods section. While we observe a significant variation in ROC AUC scores depending on the tested groups (i.e. virus families), the performance of each method is not correlated with the number of observations within each group. The *Pneumoviridae* family might be an outlier in our dataset as the predictive performance of all the considered models are substantially different for this family than those observed for the other families. Although some groups display a noticeable correlation between pHLA immunogenicity and pHLA binding affinity predictions (e.g. *Pneumoviridae* and *Orthomyxoviridae*), this trend is not confirmed across all groups. The performance (median ROC AUC across virus families) of our method is comparable to those obtained for binding affinity and ligand likelihood predictors, usually with a smaller variance of prediction performance (see Figure 5 and Figure 6).

**Figure 6.**
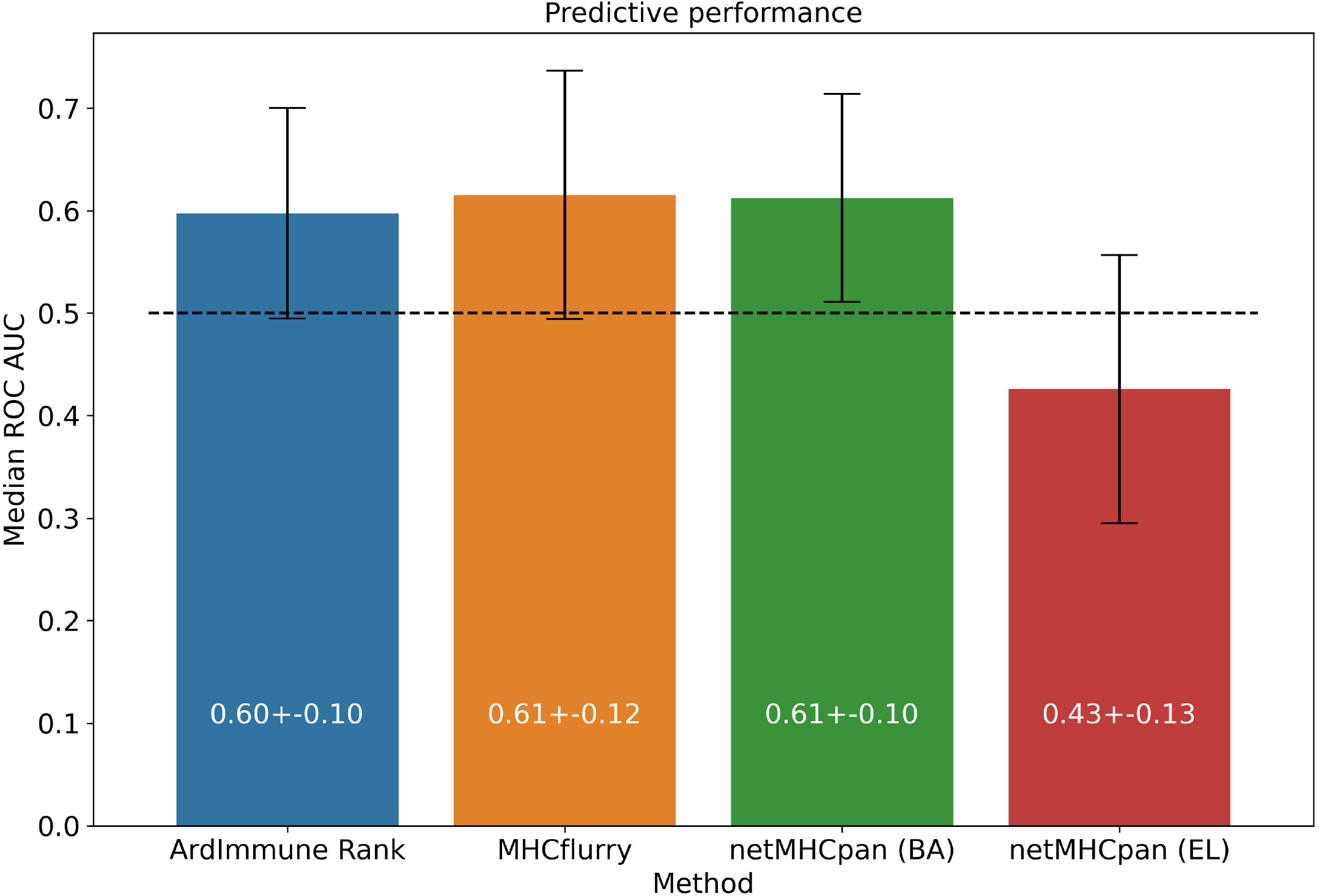
Predictive performance of the selected models, averaged across virus groups in the training dataset.

The model was then used to predict the immunogenicity of peptides from the SARS-CoV-2 proteome. Target peptides and HLA types considered for the analysis were selected according to the procedure described in the “Selection of peptides” and “Selection of HLA alleles” sections, respectively. A considerable number of peptides with high scores are observed in both structural and non-structural proteins, encompassing different HLA alleles. Structural epitopes are dominated by the Spike protein, whereas the non-structural ones mostly originate from the ORF1a/ORF1ab-encoded polyproteins. Peptides with percentile rank ≤ 2 presented across the selected HLAs, were considered for both structural (Table 3) and non-structural (Table 4) viral proteins. We noticed that some HLA alleles exhibit a large number of highly-ranked peptides, in particular A*02:01, A*11:01, A*24:41 and C*12:03. Interestingly, the presence of some of these alleles was earlier reported to be statistically correlated with the immune protection in SARS cases. Namely, A*02:01 was found to present immunogenic peptides (Ahmed et al. 2020; Lee and Koohy 2020) whereas A*11:01-restricted epitopes were proposed to be included in a SARS-CoV vaccine by Sylverster-Hvid et al. 2004. Groups of peptides predicted to be associated with multiple HLAs are shown in Figure 7. These epitopes originate from both structural and non-structural antigens.

**Figure 7.**
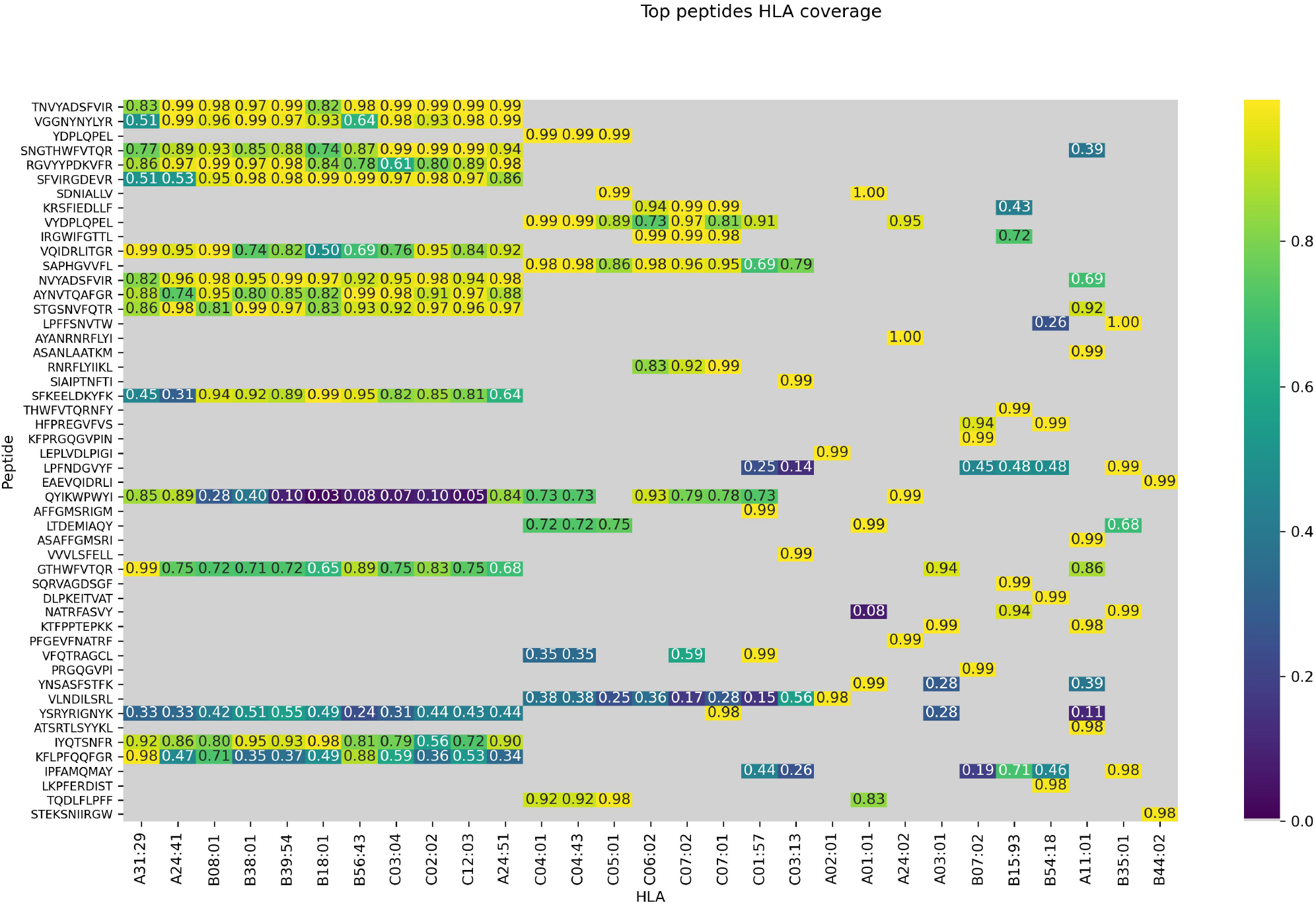
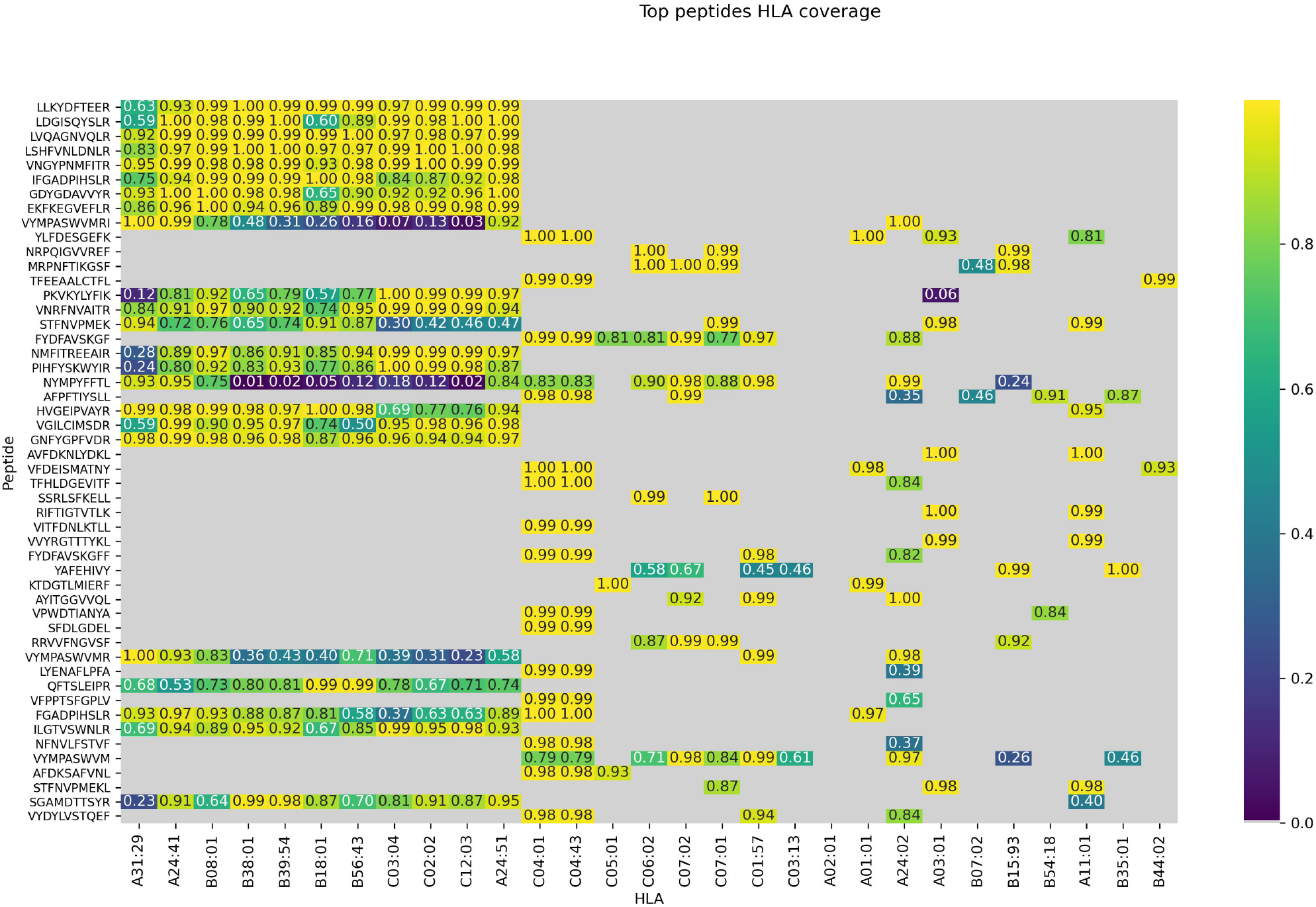
Peptides presented across multiple HLAs. Immunogenicity scores are reported for epitopes from both structural (top) and non-structural (bottom) proteins. Peptide-HLA combinations marked in grey are predicted non-binders (netMHCpan 4.0 percentile rank > 2). For the remaining pHLAs, the color relates to the percentile rank of our predictions for a given HLA type (0.95 means that the prediction is among top 5% of the predictions for that particular HLA allele).

### 3.2 SARS-CoV-2 genome diversity analysis

In order to enable the exclusion of peptides originating from genetically highly variable areas, the mutation frequency of each amino acid within the SARS-CoV-2 genome was computed (see Materials and methods for details). The genes that those peptides originate from are likely to mutate, hence the inclusion of such peptides might lower the vaccine efficacy over time. From the analysis of 8,639 complete genome sequences, obtained from different SARS-CoV-2 isolates, which then were translated into protein sequences, the mutation frequency at each amino acid position was computed. For each peptide in the SARS-CoV-2 proteome, the maximum mutation frequency was calculated (see Materials and methods), and peptides with the resulting score ≥ 0.05 (marked in color in Table 3 and Table 4) are considered to be highly variable (HV) and should be disregarded as vaccine components. 13 amino acid positions were observed to contain mutations in at least 5% of the selected sequences. Among these, as many as 9 amino acid positions were mutated in more than 10% of the selected sequences, while 2 positions showed mutations in fully half of the samples (more than 50%). In Table 5 we present the most frequently mutated positions within the SARS-CoV-2 proteome. Mutation frequency values for all positions are available in the Supplementary Data 2. Figures presenting distribution of mutation frequency are available in the Supplementary Data 3. Within the top-50 immunogenic peptides originating from the SARS-CoV-2 structural and non-structural proteins (NSPs), 1 and 3 HV peptides were found, respectively.

### 3.3 Toxicity/tolerance results

Each peptide derived from the SARS-CoV-2 proteome was studied to ascertain the lack of similarity with peptides present in the reference human proteome. When administered in a vaccine, epitopes highly similar to peptides presented by the host’s healthy tissues could either trigger an unwanted immune self-reaction or be tolerated by the immune system. In both cases, these peptides should be eliminated from the vaccine composition. A total of 11 SARS-CoV-2-derived peptides with moderate similarity to human proteins were found (E-value ≤ 4). Of these, 4 were significantly similar (E-value ≤ 1) and thus should be avoided (see Supplementary Data 1). None of these peptides were found within the top-100 ranked peptides.

### 3.4 Comparison with other methods

Results from a list of selected publications were compared with percentile ranks computed by our method for the same pHLAs. We did not find any significant correlation with the in silico predictions from Grifoni et al. 2020, Lee and Koohy 2020, and Gupta et al. 2020 highlighting a clear distinction between our methodology and the procedures used in these studies. Although the best candidate selected by Gupta et al. is not among our best candidates for HLA-A*11:01, it is scored by the model as the top candidate among those proposed by the authors. A moderate negative correlation (ρ ≅-0.45) was observed between the percentile rank scores of our method and the scores presented by Smith et al. 2020. Although our top peptide candidates associated with the HLAs proposed by Baruah and Bose 2020 do not include any of the five peptides proposed by the authors, we noticed a consensus between the HLA percentile rank of the pHLAs selected by the authors, and our percentile rank scores (Figure 8).

**Figure 8.**
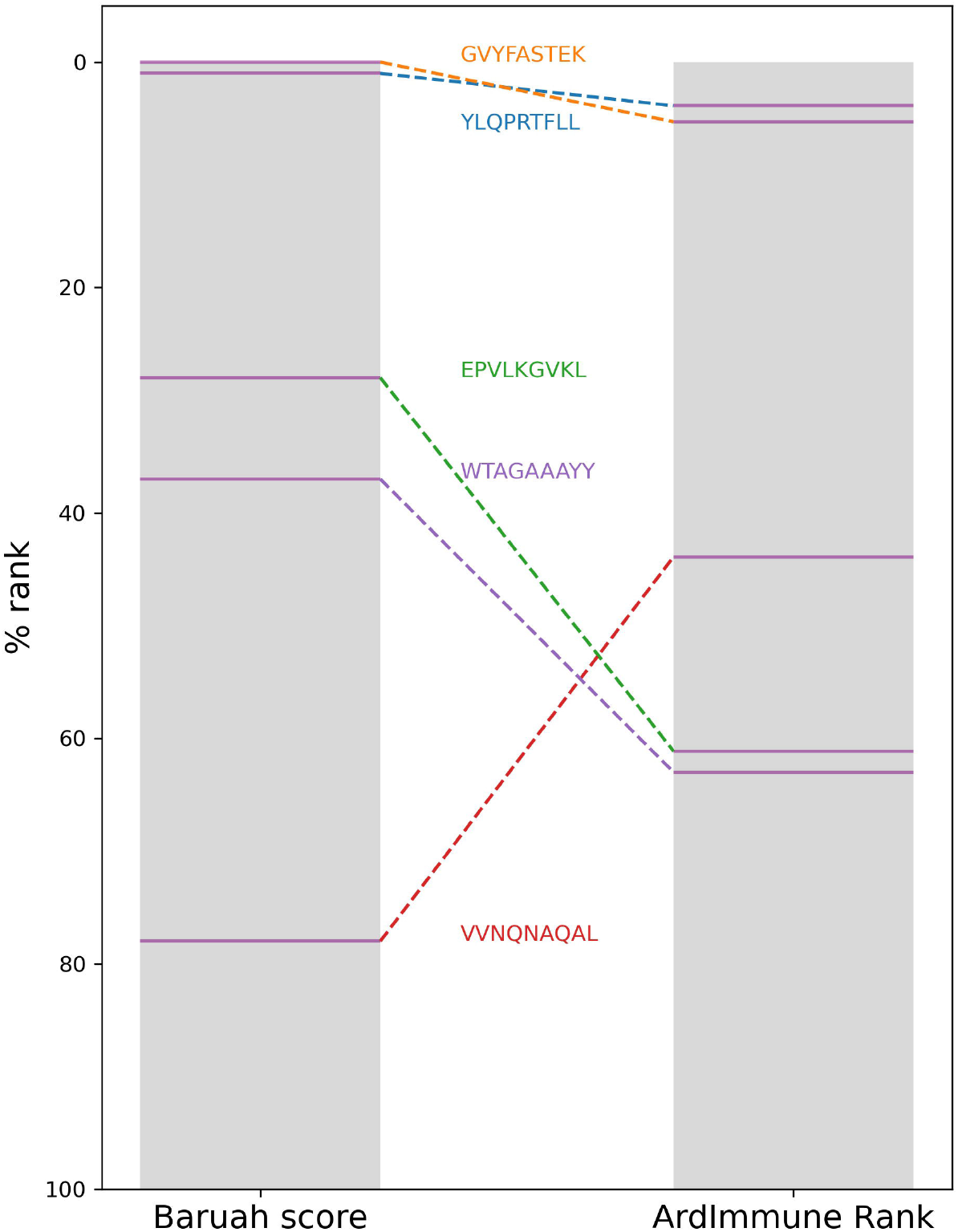
The HLA percentile ranks of the 5 peptides selected by Baruah et al. as computed from Baruah score and ArdImmune Rank.

The immunogenicity scores predicted by our model were then compared with the experimental measurement of pHLA binding stability done by Prachar et al. 2020. Peptide candidates with high immunogenicity ranks are enriched in regions with a low stability percentage (Figure 9, left; the stability percentage is defined relative to reference peptides, see Prachar et al. 2020 for details). The concordance between low immunogenicity rank and high stability percentage is more noticeable after the exclusion of peptides with low predicted binding affinity (Figure 9, right).

**Figure 9.**
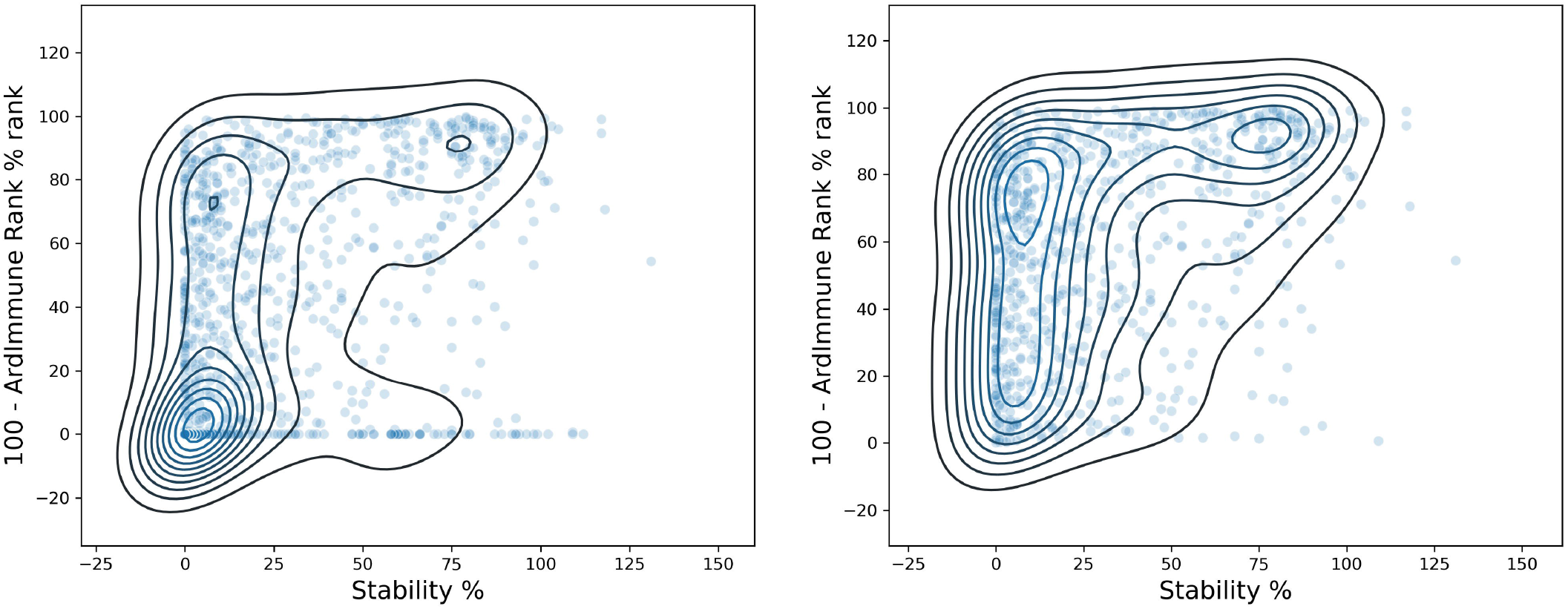
Comparison between ArdImmune Rank percentile ranks for pHLA immunogenicity and pHLA stability data measured by Prachar et al. [58] Scatter plots and kernel density estimations are shown with (right) and without (left) the exclusion of pHLA predicted non-binders (Kd percentile rank ≥ 2). The complement of the ArdImmune Rank percentile rank is shown on the y-axis (higher value = lower rank), while the stability percentage as reported by Prachar et al. is shown on the x-axis.

The Spearman correlation between pHLA stability percentage and the predicted immunogenicity (ρ = 0.392) is higher than the correlation between the stability percentage and the predicted binding affinity (ρ = 0.313). The binding affinity was computed using NetMHCpan 4.0 (Jurtz et al. 2017).

A noticeable difference in the distributions of pHLA stability percentage was obtained by ranking using binding affinity predictors and our immunogenicity predictions. A clear distinction between stable and unstable pHLAs was obtained through the selection of the top-10% and the bottom-10% scores predicted by the immunogenicity model, whereas the use of filters relying on standard binding affinity thresholds (e.g. 100nM) leads to a less defined separation (Figure 10).

**Figure 10.**
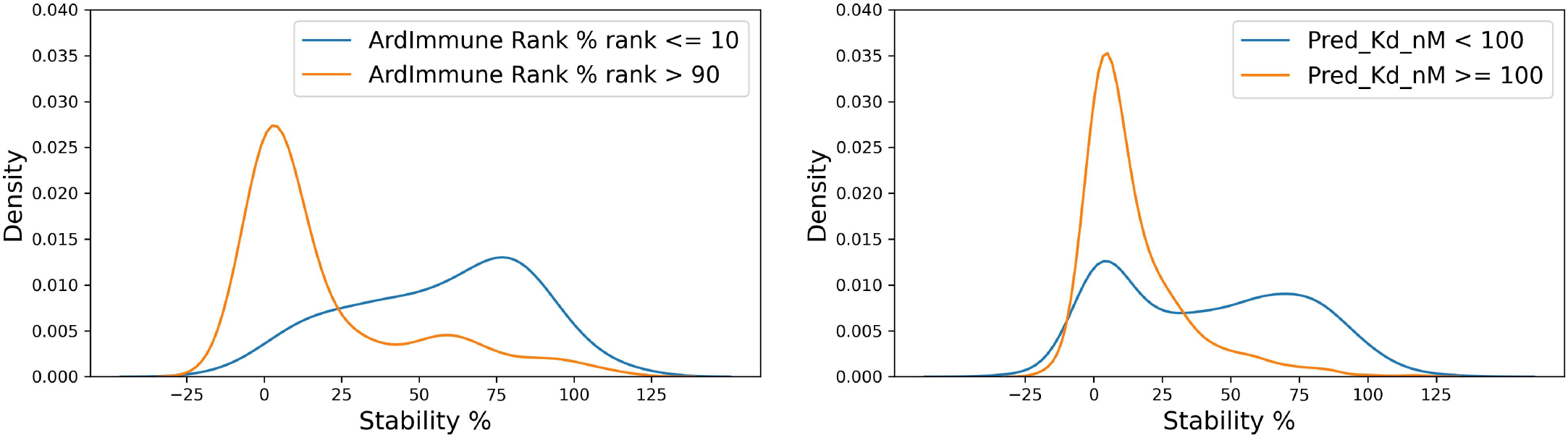
Distribution of stability percentage for different filtering procedures. The respective pHLA stability score densities of the 10% top ranked and the 10% lowest ranked peptides in terms of predicted immunogenicity is shown on the left. The pHLA stability score densities computed according to the binding affinity ranges reported by Prachar et al. [58] (K_d_ ≥ 100 nM, K_d_ < 100 nM, based on predicted binding affinity) is shown on the right.

Finally, we report low scores for all the five class I pHLAs which were experimentally confirmed to be non-immunogenic by Rammensee et al. 2020. None of these peptides was recommended by ArdImmune Rank as a candidate to be included in a vaccine formulation against SARS-CoV-2.

## 4 Discussion

The high selective pressure exerted upon coronaviruses, caused by the need of a viable host for survival, together with their high genetic variability, facilitates their rapid evolution and the prompt generation of escape mutants. Despite the vigorous effort of the industry, vaccine design, clinical trials, and production require at least several months and most likely several years. Many investigations aimed at developing vaccines protecting humans and animals from coronaviruses were initiated in the last few decades, setting the basis for the recent scientific advancement in COVID-19 treatment. Nonetheless, a limiting aspect associated with the approval and commercialization of a vaccine is that the demand for a vaccine is limited to the outbreak period, and its market value is proportional to the number of people affected. This represented a major issue for the development of vaccines for SARS and MERS (Dhama et al. 2020; Du et al. 2009). In addition, the majority of coronavirus biotherapeutics (i.e., antibodies and vaccines) are designed to leverage neutralizing antibodies directed against the S protein. Safety issues such as those associated with the ADE and CSS events, make the development of vaccine and antibody-based therapies even more problematic. In combination with the stimulation of humoral immune response, which is aimed at the direct neutralization of the virus, the targeted elimination of infected cells is a crucial element of the immune response against viruses. This might be induced either by the administration of a vaccine eliciting protective CD8+ Cytotoxic T Lymphocyte (CTL) or by transferring CD8+ cells engineered to recognize viral antigens specifically. Previous studies have confirmed a strong correlation between the depletion and exhaustion of T-cells and worse prognosis in critical coronavirus patients (Diao et al. 2020) highlighting the potential of vaccines inducing T-cell responses for COVID-19 prevention. This strategy has beneficial features such as a lower risk of stimulating ADE and CSS with respect to antibody-based strategies (Channappanavar et al. 2016; Jaume et al. 2011) and the stimulation of the immune response against intracellular epitopes not reachable by the antibodies but potentially highly immunogenic. In both cases, the selection of effective immunogenic epitopes is of paramount importance.

The aim of this study was to identify SARS-CoV-2 epitopes for the development of a vaccine composition focused on T-cell activation. We investigated several aspects pre-determining whether viral epitopes may induce an effective T-cell response, including the MHC-I peptide presentation and immunogenicity potential, SARS-CoV-2 genome variability, and possible toxicity/immune tolerance of the peptides considered.

In contrast to the majority of works on this topic either relying of pHLA binding and presentation events or modeling single pHLA structural interactions, the model applied herein was designed to leverage simultaneously information about the propensity of a peptide to be presented by its cognated HLA and the probability that such pHLA is immunogenic, inferred from similar experimental data. As we show in Figure 3 when evaluated on the experimentally-validated *Coronaviridae* immunogenicity data, our approach has higher performance than the widely-used binding affinity and pHLA presentation predictors.

By applying our method, a considerable amount of highly scored T-cell epitopes was found across the SARS-CoV-2 proteome, encompassing the structural proteins and NSPs, as shown in Tables 3 and 4. The majority of selected epitopes were conserved across different SARS-CoV-2 isolates. Only 16 epitopes were excluded because of their significant mutability (see Table 5). The availability of epitopes from NSPs allows for the design of vaccine components dedicated to T-cell responses, and might be further integrated with other components focused on B-cell responses. The adoption of such a compartmentalized strategy might help to lower the risk of non-neutralizing antibody production, which constituted a reason of concern during the development of a vaccine formulation for SARS. Moreover, during the early stages of viral infection, the expression of non-structural proteins is significantly higher than the expression of structural ones. The targeted stimulation of the immune response towards epitopes originating from non-structural proteins might be useful to induce an immune response at the early phase of the disease. Some highly ranked peptides were found to be presented across multiple HLAs and could be used to increase population coverage while decreasing the number of epitopes needed to be included in the vaccine formulation. This aspect could be particularly relevant for solutions relying on delivery systems of limited capacity.

The risk of eliciting potentially harmful and sometimes deadly (Linette et al. 2013) cross-reactivities is an issue to be carefully addressed in vaccine design. On the other hand, epitopes shared with proteins from the host could also be tolerated by the host’s immune system, being not useful for vaccine purposes. Considering the importance of such an aspect, the analysis of potential toxicity and tolerance was addressed in this study, leading to the identification of 4 highly ranked epitopes having a certain degree of similarity with proteins within the human proteome. Such peptides were removed for safety and efficacy reasons.

The substantial difference between the selection of pHLA candidates performed by our methodology with respect to those presented by Grifoni et al. 2020, Lee and Koohy 2020 and Gupta et al. 2020 highlights a clear distinction between these approaches. Nonetheless, our method supported the selection of top candidates in small datasets obtained by applying hand-crafted filtering stages (Baruah and Bose 2020; Gupta et al. 2020). The mild correlation with the results from Smith et al. 2020 might indicate the usage of equivalent components during some steps of the selection process. A relative concordance between the pHLA stability scores from Prachar et al. 2020 and the associated immunogenic scores computed by our method was observed (Figure 9). Moreover, we show that the peptide ranks produced by our immunogenicity model have a higher correlation with the experimentally measured pHLA stability than the ranks obtained by methods relying solely on binding affinity or ligand likelihood predictions. This observation is consistent with works reported in the literature (Harndahl et al. 2012). We also obtained low immunogenicity scores for all five peptides which have been experimentally confirmed by Rammensee to be unable to activate CD8+ lymphocytes.

## 5 Conclusions

In this paper we suggested a SARS-CoV-2 vaccine composition in the form of the list of epitopes optimized for their (predicted) immunogenicity and HLA population coverage. Our motivation is that cellular immune response is fundamental for an effective SARS-CoV-2 vaccine and it also mitigates the risks of ADE and CSS which are typically associated with modalities relying on the activation of humoral immune response. We showed that the predictive model, on which our methodology is based outperforms, on *Coronaviridae* data, other methods used to date for the design of epitope-based vaccines against SARS-CoV-2. Our approach differs from other existing methods and shows an improved consistency with experimental data. This includes a higher correlation with the measured pHLA stability in comparison with methods based solely on binding affinity predictions. The limitations of our method have the same roots as those found in other in-silico approaches based on predicting various pHLA properties, i.e. the accuracy of these predictive methods. We expect that with the increasing amount of experimentally validated data and with further algorithmic enhancements in the field of artificial intelligence, the accuracy of such models and the effectiveness of vaccine design will continue to improve. In spite of these limitations, the combination of genomic analysis and AI techniques already represents a viable methodology for the rational design of epitope-based vaccine formulations for COVID-19 prevention.

## 6 Authors’ contributions

GM wrote the article with contributions from IN, PSk and JK. AM, PSk and IN performed the analyses and generated figures and tables included in the article. GM, IN, AM, PSk, JK, ASD, KG, PK, MD developed the applied methodology. PSt conceived the idea for the project and coordinated the work. ASD, MS and KP gave essential contributions to the interpretation of immunological and virological aspects of the study. All the authors reviewed, edited, contributed to the article and approved the submitted version.

### 7 Acknowledgments

Ardigen and COVID-19 Vaccine Corporation (CVC) announced that they entered a research collaboration aimed at the development of SARS-CoV-2 vaccine.

## 10 Data availability statement

The lists containing the predicted immunogenic peptides with percentage rank ≤ 2 are included in this study (Supplementary Tables 3 and 4). The list of all the predicted immunogenic peptides generated during this study are available from the corresponding author upon reasonable request.

