## Supplementary figures and images for "AI aided design of epitope-based vaccine for the induction of cellular immune responses against SARS-CoV-2"

### Supplementary Data 3

#### Supplementary Materials 1

Mutation frequency for each position within all SARS-CoV-2 proteins.

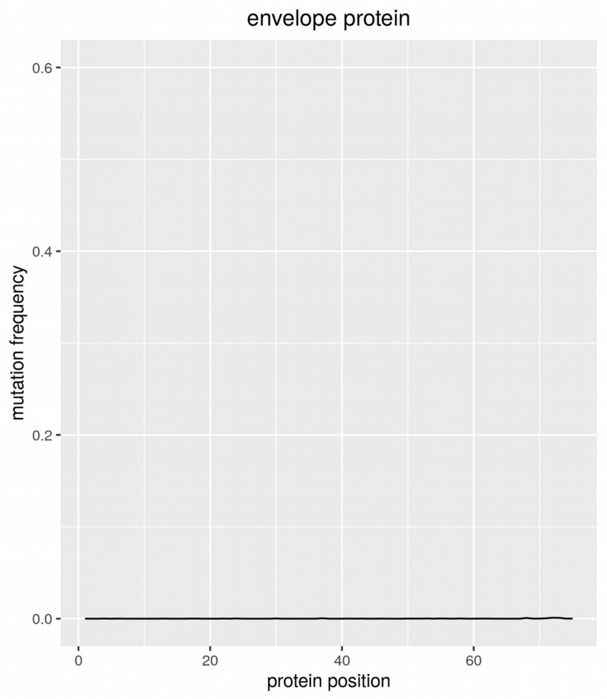


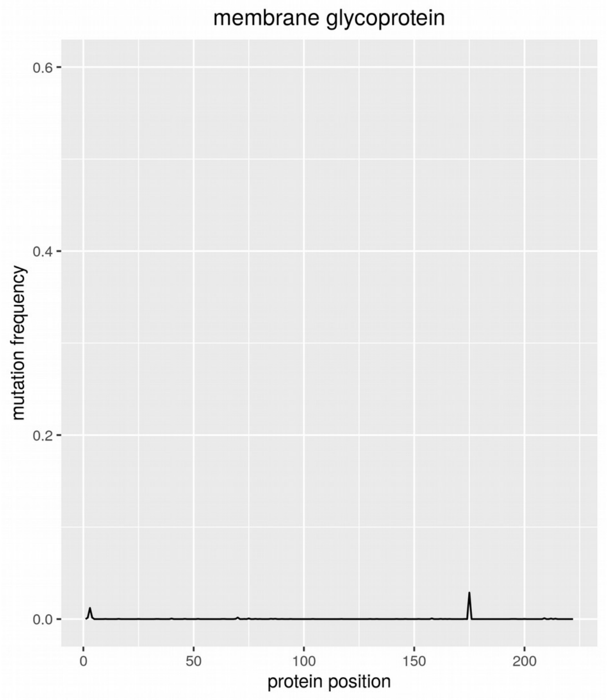


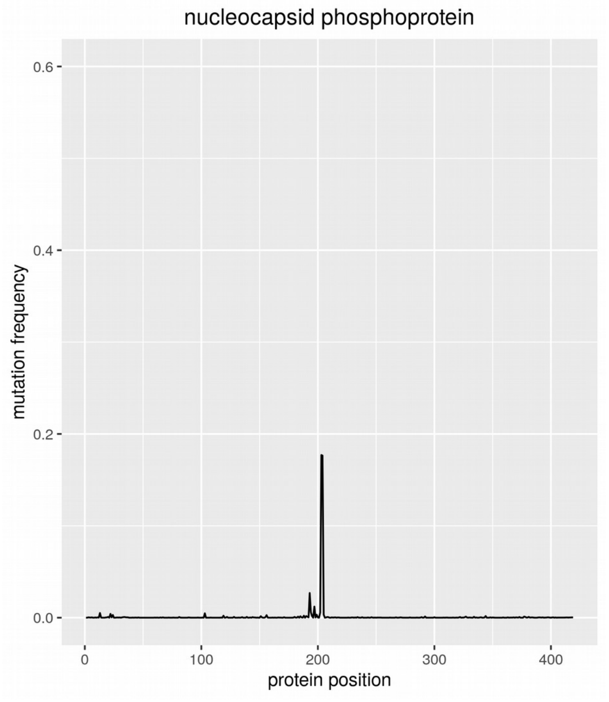


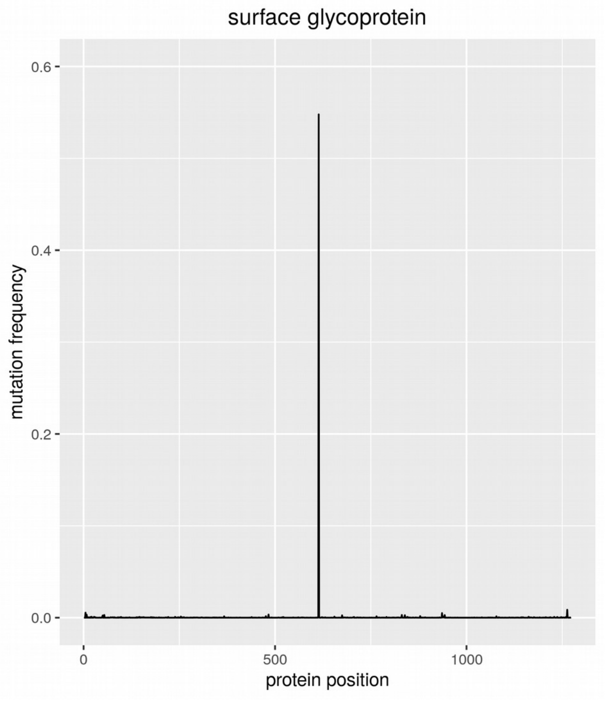

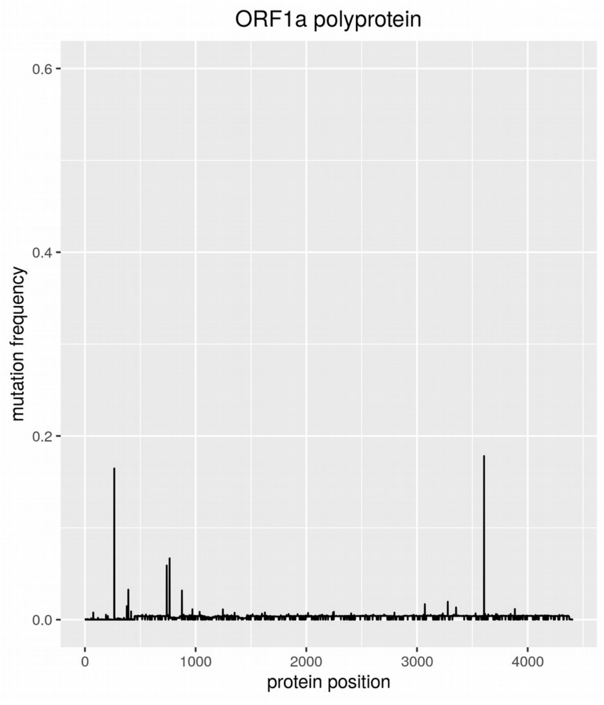

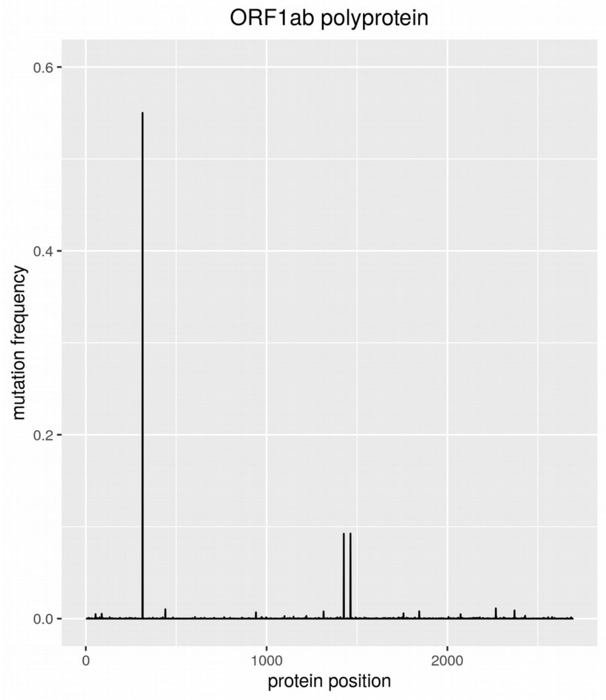

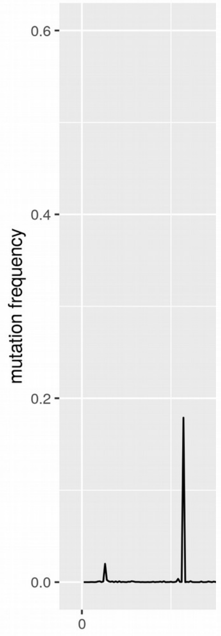

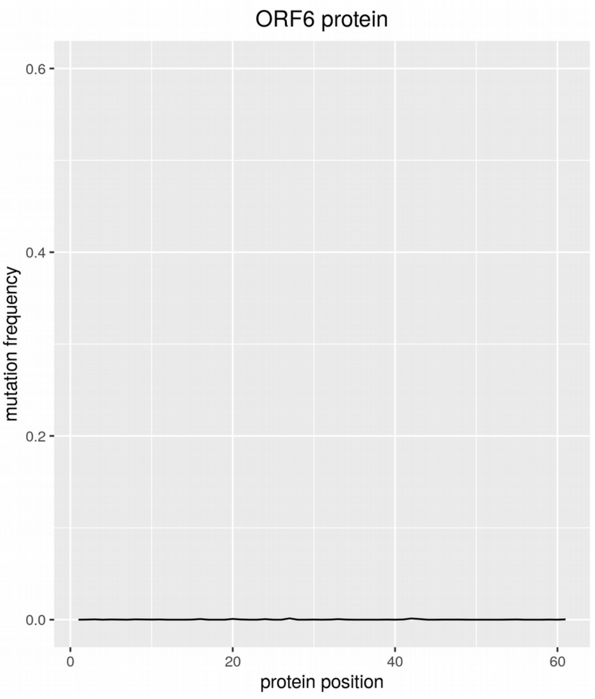

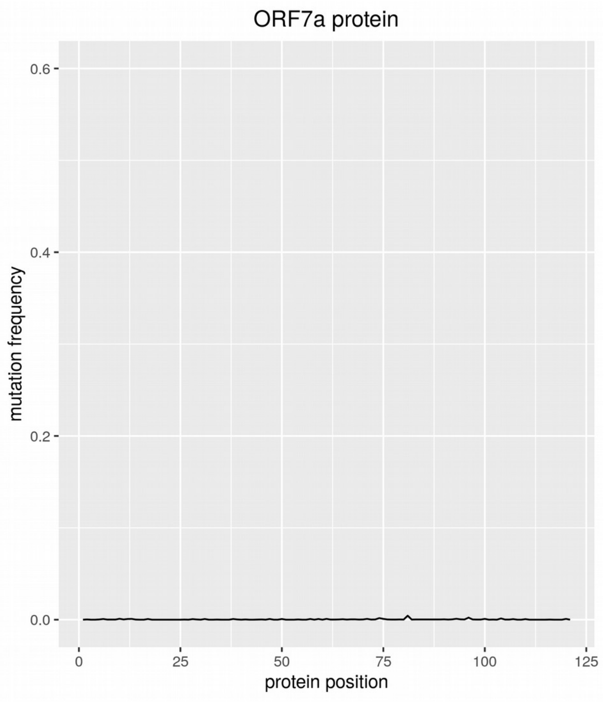

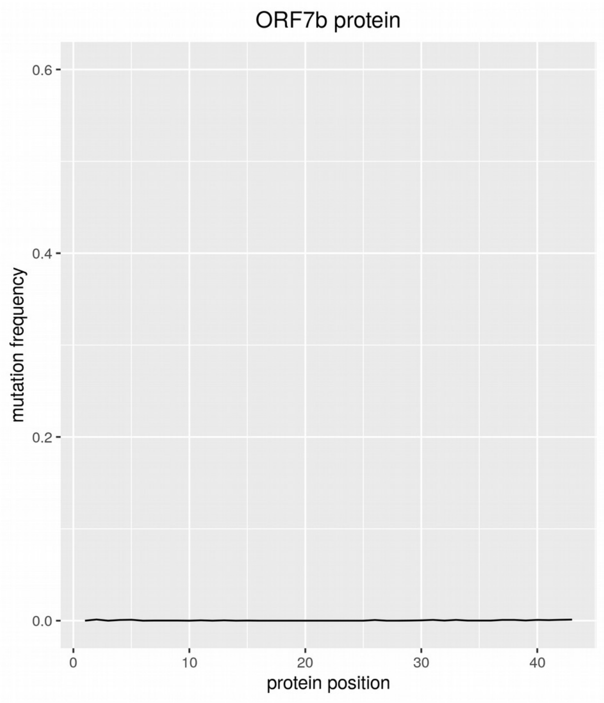

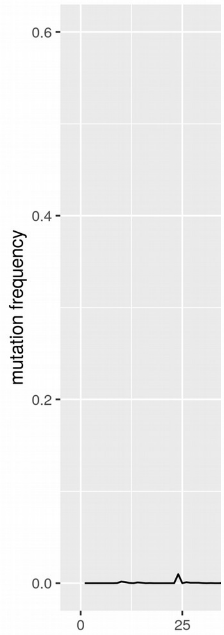

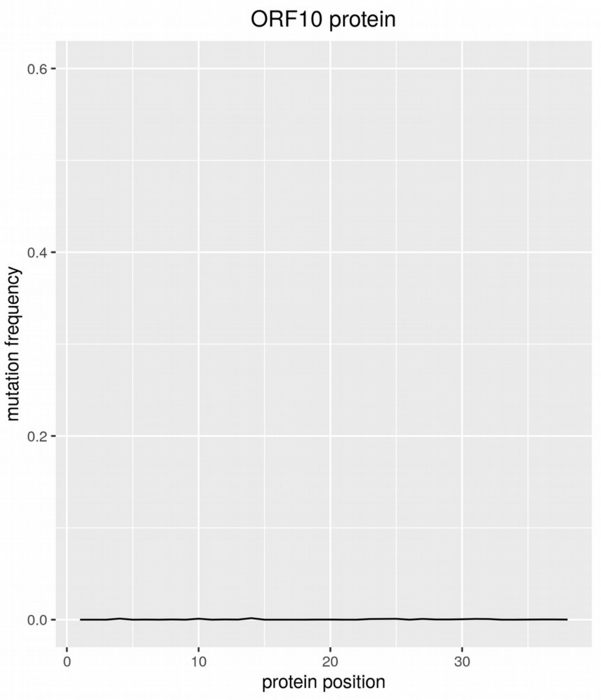
